# A phage satellite manipulates the viral DNA packaging motor to inhibit phage and promote satellite spread

**DOI:** 10.1101/2024.04.22.590561

**Authors:** Caroline M. Boyd, Kimberley D. Seed

## Abstract

ICP1, a lytic bacteriophage of *Vibrio cholerae*, is parasitized by phage satellites, PLEs, which hijack ICP1 proteins for their own horizontal spread. PLEs’ dependence on ICP1’s DNA replication machinery, and virion components results in inhibition of ICP1’s lifecycle. PLEs’ are expected to depend on ICP1 factors for genome packaging, but the mechanism(s) PLEs use to inhibit ICP1 genome packaging is currently unknown. Here, we identify and characterize Gpi, PLE’s indiscriminate genome packaging inhibitor. Gpi binds to ICP1’s large terminase (TerL), the packaging motor, and blocks genome packaging. To overcome Gpi’s negative effect on TerL, a component PLE also requires, PLE uses two genome packaging specifiers, GpsA and GpsB, that specifically allow packaging of PLE genomes. Surprisingly, PLE also uses mimicry of ICP1’s *pac* site as a backup strategy to ensure genome packaging. PLE’s *pac* site mimicry, however, is only sufficient if PLE can inhibit ICP1 at other stages of its lifecycle, suggesting an advantage to maintaining Gpi, GpsA, and GpsB. Collectively, these results provide mechanistic insights into another stage of ICP1’s lifecycle that is inhibited by PLE, which is currently the most inhibitory of the documented phage satellites. More broadly, Gpi represents the first satellite-encoded inhibitor of a phage TerL.

## INTRODUCTION

Bacteriophages (phages) are highly abundant viruses that infect and hijack bacteria to produce progeny phages. During an infection, some phages can activate resident, integrated phage satellites. These small (<20 kb) satellites follow a similar lytic lifecycle as their phages. They replicate and then package their genomes into virions, which are released from the cell and can spread horizontally to neighboring cells. Satellites, unlike phages, lack some essential proteins required to produce these virions and thus are reliant on those encoded by their “host” or “helper” phage (reviewed in 1). Due to the thievery of essential proteins by the satellite from the host phage, there is typically competition between phage and satellite for these resources. Often, satellites outcompete their host phages, and this leads to an antagonistic relationship wherein satellites promote the production of their progeny at the cost of the host phage. Many satellites are dependent on their host phage’s proteins for genome packaging. In the case of most double-stranded DNA phages, genome packaging is regulated by a terminase complex. A specificity factor, the small terminase (TerS) recognizes phage DNA, and a motor complex, the large terminase (TerL), coordinates cleavage and packaging of the genome into preformed capsids (called procapsids) (reviewed in 2). TerS binds to a unique sequence (referred to as *cos* or *pac* site) on the phage genome to initiate packaging by recruiting TerL to the DNA at this site. DNA is then forcefully packaged into the procapsid by the TerL motor. Packaging terminates after precisely (for *cos* site directed packaging) or slightly more than (for *pac* site directed packaging) one unit length of the genome is packaged within the capsid, triggering TerL to cleave the DNA a second time (reviewed in 3).

Most satellites lack their own packaging motor and depend on the TerL from their host phage (1). Various satellites that rely on the phage TerL have evolved strategies to compete with the phage for the use of this protein to package their genomes. Briefly, different satellites have been described to 1) disguise their genomes to look like the phage genome by encoding a highly similar packaging sequence as their host phage (4–6), 2) encode a protein that binds to and modifies the phage TerS to redirect the complex by increasing its binding affinity to a similar, but distinct sequence on the satellite (7), or 3) encode an alternative TerS which recognizes the unique satellite packaging signal while simultaneously deploying a protein that directly interferes with the function of the phage’s TerS, thus inhibiting phage genome packaging (8, 9). All these strategies promote satellite packaging, while the latter two strategies also directly antagonize phage packaging. Satellite-directed inhibition of phage genome packaging in these systems is not completely inhibitory to phage production, as all three scenarios result in both satellite and phage progeny production.

Satellites can be grouped into families, which are defined by their genetic similarity. Satellites from different families do not have similar genetic sequences, suggesting that rather than originating from a single common ancestor, satellites evolved in parallel. Interestingly, many satellites have converged on parasitizing their host phages at the same stages of the lytic cycle (1). However, independent paths of evolution allowed for innovation in mechanisms that achieve the same result, as described above for satellite genome packaging. PLEs are a family of satellites that parasitize the *Vibrio cholerae* phage, ICP1, and are the only satellites currently known that fully inhibit progeny production of their host phage (10, 11). To date, 10 PLEs have been identified share their genetic organization and a subset of, but variants are differentiated by unique regions found only in that variant (28). PLEs are known to inhibit ICP1’s capsid production (12), genome replication, and genome packaging (13). Mechanistically, PLEs use TcaP, an external scaffolding protein, to redirect ICP1 coat proteins to form smaller capsids that do not accommodate the full length of the ICP1 genome (12). PLEs also deploy NixI, a nicking endonuclease, to target ICP1’s genome and reduce ICP1 replication (13). Neither TcaP nor NixI is necessary for PLEs to block ICP1 progeny production (14), and, interestingly, NixI is insufficient to explain PLEs’ interference with ICP1 specifically at the stage of genome packaging (13). These results suggest that PLEs use an unknown mechanism(s) to interfere with ICP1 genome packaging. We hypothesized that PLEs encode a dedicated inhibitor of ICP1’s packaging machinery (Figure 1A). Given the precedent in other satellites, we hypothesized that PLEs would also encode a mechanism to promote their own packaging (Figure 1A).

**Figure 1.**
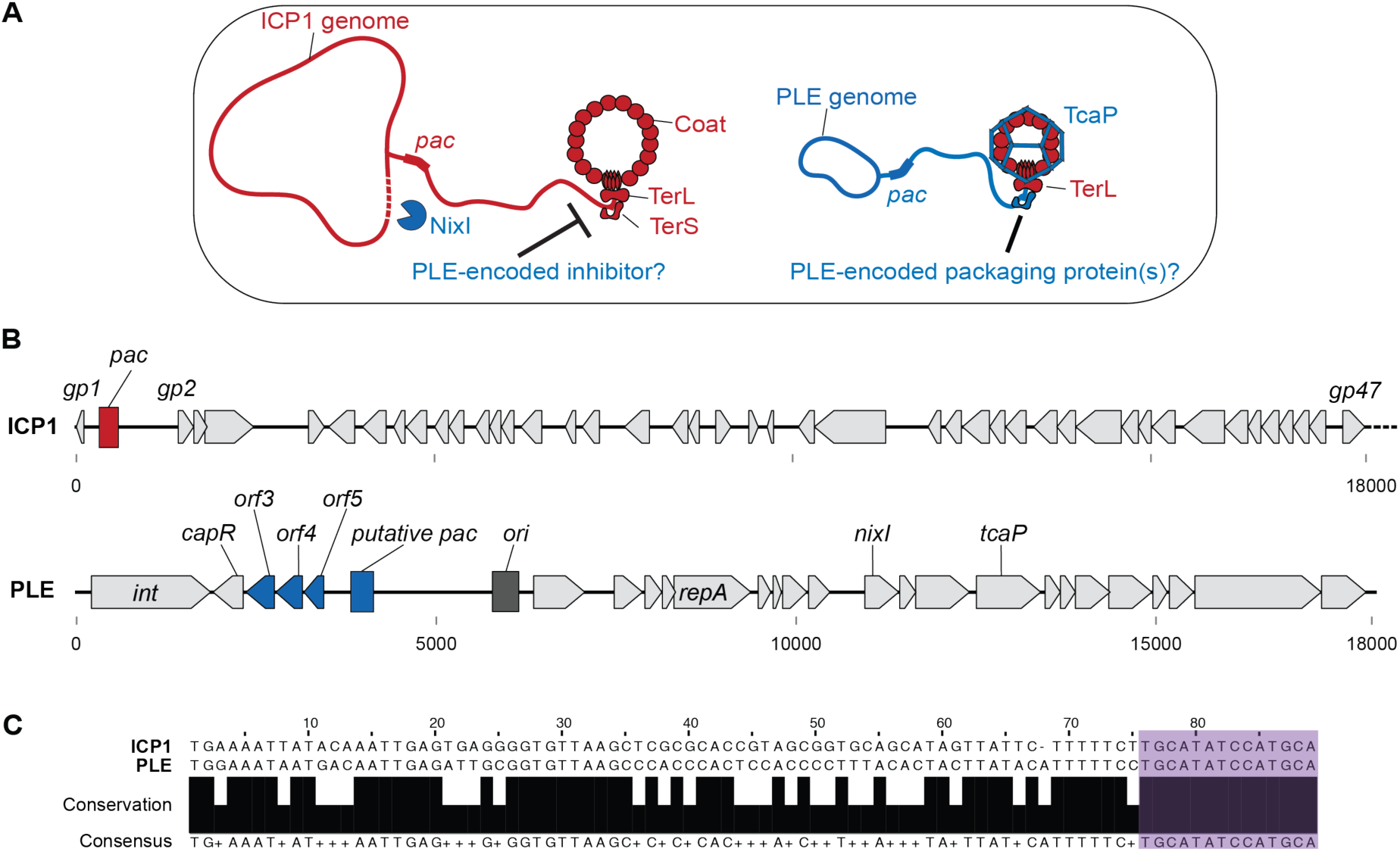
Identification of PLE1’s putative *pac* site and packaging proteins. (A) Model for hypothesized ICP1 and PLE1 genome packaging strategies wherein ICP1 and PLE depend on the ICP1-encoded motor complex, TerL, to fill procapsids. Presumably, ICP1 relies on a TerS to recognize its genome. In addition to PLE1 encoding an inhibitory nicking endonuclease that targets ICP1’s genome, NixI, and an external scaffold that directs assembly of small capsids, TcaP, PLE1 likely encodes an ICP1 genome packaging inhibitor as well as proteins to promote PLE1 genome packaging. DNA and proteins encoded by ICP1 and PLE1 are shown in red and blue, respectively. (B) Gene maps of a fragment of ICP1’s genome and the entirety of PLE1’s genome show the location of the putative *pac* sites as red and blue boxes on ICP1 and PLE1 genomes, respectively. PLE1’s origin of replication (*ori*) is shown as a gray box. Genes in PLE1 relevant to this work are labeled, and genes of interest for their potential roles in packaging are colored blue. The genomes are to scale, marked by 5 kb tick marks. (C) Sequence alignment of the BLAST hit of PLE1’s genome against ICP1’s showing similarity between ICP1’s *pac* site and PLE1’s putative *pac* site. The identical 14 base pairs between PLE1 and ICP1 in this region are highlighted in purple.

In this work, we set out to define PLE’s strategy to preferentially package its genome. Our work reveals that PLE uses a combination of three previously described packaging strategies with some unique twists. First, PLE partially mimics ICP1’s *pac* site. PLE encodes a 14-base pair signal within an 89-base pair sequence that matches part of ICP1’s *pac* site and is required for PLE transduction. Second, PLE produces an inhibitor that blocks ICP1 genome packaging. PLE’s Orf3 (Gpi) directly binds to ICP1’s TerL, rather than the phage’s TerS, as is seen in other satellites, and indiscriminately blocks genome packaging. Third, and in line with PLE’s dependence on ICP1’s TerL for its own packaging, PLE requires two additional proteins, Orf5 (GpsA) and Orf4 (GpsB), to overcome Gpi’s negative effect on genome packaging. Genetic data show that GpsA and GpsB specifically contribute to PLE genome packaging in the presence of Gpi, while ICP1’s genome packaging remains inhibited. Gpi, GpsA, and GpsB individually contribute to PLE packaging and, together with the PLE *pac* site, are sufficient for high-frequency transduction. Curiously, PLE can achieve moderate levels of packaging in the absence of its three packaging proteins, but other PLE-encoded factors are required to antagonize ICP1 under these conditions. Functional conservation of PLE’s packaging genes in all known PLE genomes supports the advantage of increased frequency of packaging such that PLE does not depend only on *pac* site mimicry. This work provides the initial characterization of PLE’s genome packaging strategy and reveals a previously undescribed combination of *pac* site mimicry and packaging redirection. Additionally, this work provides the first example of an inhibitory satellite protein (Gpi) that directly binds to a phage-encoded TerL to block genome packaging.

## MATERIAL AND METHODS

### Bacterial Growth Conditions

*V. cholerae* and *E. coli* were grown with aeration at 37°C on LB agar or in LB broth. Where needed, antibiotics were used at the following concentrations: streptomycin 100 µg/mL, kanamycin 75 µg/mL, spectinomycin 100 µg/mL, carbenicillin 50 µg/mL, trimethoprim 32 ug/mL, chloramphenicol 2.5 µg/mL (solid media) or 1.25 µg/mL (liquid media) for *V. cholerae* and 25 µg/mL for *E. coli*.

### Genetic Manipulations

#### Plasmid Construction

Plasmids were constructed by Gibson Assembly or Golden Gate Assembly and then purified in *Escherichia coli* XL1-Blue. Plasmids were electroporated into *E. coli* BTH101 for bacterial adenylate cyclase two-hybrid (BACTH) assays or into *E. coli* S17s for mating into *V. cholerae*.

#### Transformation of V. cholerae

Strains with PLE1 marked with a kanamycin resistance cassette downstream of the last ORF (10) were made naturally competent by methods previously described (15) and then were transformed with DNA fragments generated from purified PCR products. Products contained either a spectinomycin resistance marker flanked by FRT recombinase sites assembled to up and downstream regions of homology by splicing by overlap extension PCR as previously described (15) or a carbenicillin resistance marker flanked by regions of homology. Spectinomycin-resistant transformants were transformed with a plasmid carrying a FLP recombinase, which was induced by 1 mM isopropyl-β-D-thiogalactopyranoside (IPTG) and 1.5 mM theophylline, allowing for removal of the spectinomycin resistance cassette via recombination, resulting in an in-frame FRT-marked deletion. The strains were then cured of the plasmid.

#### ICP1 genome editing

Specific editing of ICP1’s genome was carried out by CRISPR-Cas engineering as described previously (16). Briefly, a strain of *V. cholerae* with an inducible CRISPR-Cas system was engineered with a plasmid encoding spacers targeting ICP1 and a repair template that provided escape from CRISPR targeting. The *V. cholerae* host was grown to mid-log, induced with 1 mM IPTG, and then infected with ICP1. Plaques were purified on a targeting-only strain and sequence verified.

All deletions, insertions, and plasmid constructs were confirmed by PCR and Sanger sequencing. A complete list of strains used in this study can be found in Table S1.

### ICP1 plaque assays

*V. cholerae* overnight cultures were back diluted to OD_600_=0.05 and grown with aeration in LB supplemented with chloramphenicol (where appropriate) at 37°C to OD_600_=0.2 then induced with 1 mM IPTG and 1.5 mM theophylline (where appropriate) for 20 minutes to reach OD_600_=0.3, then mixed with pre-diluted phage samples. A 7-10 minute incubation allowed for phage attachment before the samples were plated in 0.5% molten top agar (supplemented with antibiotics and inducer where appropriate) and then incubated overnight at 37°C.

### ICP1 mutant purification and whole genome sequencing

*V. cholerae* expressing Orf3 from a plasmid was used in a plaque assay as described above with ICP1_2018_Mat_B. Rare plaques were picked and purified three times on the same host and then high titer stocks were generated. Non-encapsidated DNA was degraded by treating phage stocks with DNase for 30 minutes at 37°C, and then the enzyme was heat-inactivated. Capsid-bound phage genomic DNA was isolated using a Qiagen DNeasy blood and tissue DNA purification kit (Qiagen, 69506) according to the manufacturer’s protocols. Illumina sequencing was performed by the Microbial Genome Sequencing Center (Pittsburgh, PA) at a 200Mb depth. The genomes were assembled using SPAdes and then analyzed by BreSeq (v0.33).

### Bacterial adenylate cyclase two-hybrid (BACTH) assays

Genes of interest were cloned into plasmids encoding the T-25-N, T-25-C, T-18-N, or T18-C fragments, including a linker by Gibson Assembly (17). Plasmids were double transformed into electrocompetent BTH-101 cells and selected on kanamycin and carbenicillin LB agar plates. Individual colonies were picked into 200 µL of LB supplemented with kanamycin and carbenicillin and grown for 4-6 hours at 37°C with aeration. 10 µL were spotted onto LB agar plates supplemented with kanamycin, carbenicillin, 1 mM IPTG, and 40 ug/mL 5-bromo-4-chloro-3-indolyl β-D-galactopyranoside (X-Gal), allowed to air dry, and incubated at 30°C overnight.

### Post-infection deep sequencing and coverage mapping

*V. cholerae* carrying an empty vector or a vector encoding *orf3* were grown and induced as described above. 2 mL of cells were infected with ICP1 at a multiplicity of infection (MOI) of 1. 30 minutes post-infection 1 mL of culture was collected in 1 mL ice-cold methanol to halt DNA synthesis, pelleted and washed twice in ice-cold PBS. Total DNA was isolated using a Qiagen DNeasy blood and tissue DNA purification kit (Qiagen, 69506). Briefly, the samples were treated as follows: addition of 180 µL ATL and 40 µL proteinase K, incubation for 20 minutes at 56°C, addition of 4µL RNase A, 400 µL AL, and 400 µL ethanol, then bound to the column, washed twice, and eluted in ddH_2_O. Illumina sequencing was performed by SeqCenter (Pittsburgh, PA) at a 200Mb depth. Reads were mapped to reference sequences, and average coverage was calculated from three biological replicates. Mapped reads distribution for each genomic element was calculated as the percent of the total mapped reads across each element (*V cholerae* chromosome I and II, ICP1, and the plasmid) normalized by element length.

### Virion production for transmission electron microscopy

#### Virion production

50 mL cultures of *V. cholerae* strains were grown and induced (where appropriate) as described above. Lysates were collected following ICP1 infection at an MOI of 2.5. Debris was removed by a 10-minute centrifugation step at 5,000 x *g,* and then the particles were concentrated by centrifugation at 26,000 x *g* for 90 minutes. Particle pellets were nutated in Phage Buffer 2.0 (50 mM Tris-HCl, 100 mM NaCl, 10 mM MgSO_4_, 1 mM CaCl_2_) overnight, then mixed with 1:1 chloroform for 15 minutes. Centrifugation at 5,000 x *g* for 15 minutes removed chloroform and any remaining bacterial debris. The aqueous layer containing the particles was collected.

#### Transmission electron microscopy

Copper mesh grids (Formvar/Carbon 300, Electron Microscopy Sciences) were loaded with 5 µL samples for 60 seconds, wicked, immediately washed with sterile ddH_2_O for 15 seconds, wicked, immediately stained with 1% uranyl acetate (Electron Microscopy Sciences, 22400-1) for 30 seconds, wicked and completely dried. A FEI Tecnai-12 electron microscope operating at 120 kV was used to collect micrographs.

### Transduction assays

#### PLE transductions

PLE transduction assays were performed as previously described (10, 18). Briefly, PLE1 was marked with a kanamycin resistance cassette downstream of the last ORF, infected with ICP1 at an MOI of 2.5 for 5 minutes. Unbound phage were removed by 5,000 x *g* centrifugation for 2 minutes, a wash in 1 mL LB, then infected cells were resuspended (with inducer where appropriate) and incubated at 37°C for 20 minutes. The resulting lysates were treated with 10 µL chloroform. Centrifugation at 5,000 x *g* for 15 minutes removed the chloroform and bacterial debris. 20 µL of the supernatant were added to 180 µL of saturated overnight culture of recipient *V. cholerae* cells (Δ*lacZ::spec^R^*), which was supplemented with 10 mM MgSO_4_ immediately prior to transduction. The mixtures were incubated for 20 minutes at 37°C with shaking (220 rpm) and then diluted and plated on LB agar plates supplemented with spectinomycin and kanamycin. Here, one colony represents one PLE virion or transducing unit.

#### ComPacT-PLE transductions

ComPacT-PLE transduction assays were similar to PLE transduction with the following modifications: *repA* (and where appropriate, plasmid-based gene expression) were induced 20 minutes prior to infection with 1mM IPTG and 1.5 mM theophylline, ComPacT-PLE was marked with a spectinomycin resistance cassette downstream of the integrase, and unmarked or trimethoprim resistant (Δ*lacZ::trim^R^*) recipients were used, and appropriate selection was used in the LB agar plates. As a control, lysates were plated to confirm the lack of donor cell carryover in this assay. Lysates were also used for plaque assays (as described above) to quantify ICP1 production.

## RESULTS

### PLE encodes a sequence similar to ICP1’s *pac* site

PLEs are potent inhibitors of their host phage, ICP1, yet the complete suite of mechanisms controlling this inhibition remains to be elucidated. To date, it is known that TcaP restricts complete packaging of the ICP1 genome by modifying the size of the capsids (12) and that NixI restricts ICP1 genome replication by targeting and nicking the genome (13). Interestingly, ICP1 packaging is still inhibited in the absence of NixI, as indicated by the lack of cleavage around the *pac* site late during infection (13). To identify how PLE inhibits ICP1 genome packaging and to probe how PLE promotes its own genome packaging, we considered that some satellites mimic their host phage packaging signals to use the phage-encoded terminase complex for genome packaging (4–6). To see if PLE uses mimicry, we analyzed the genome of PLE1 (referred to as PLE or simplicity where possible) for similarity to ICP1’s, even though phage satellites typically lack sequence relatedness with their host phages (1). Previous work identified a 489 base pair intergenic sequence between *gp1* and *gp2* as the sequence that contains ICP1’s predicted *pac* site (19) (Figure 1B). Intriguingly, BLASTN analysis revealed an 89-base pair fragment of PLE that has 71% identity to an 88-base pair region within the putative ICP1 *pac* site (Figure 1C). Internally, this fragment contains a 14-base pair sequence that is 100% identical to part of ICP1’s *pac* site. The only other region with significant similarity was the gene encoding PLE1’s capsid operon repressing protein, CapR, which matched to a homing endonuclease gene, Gp165. The similarity of these proteins has previously been described (18) and is potentially the result of an ancient homing and domestication event (20). Other hits comprise nonsignificant (E-values greater than 0.1) regions of 15-36 base pairs (Table S2), demonstrating the expected lack of sequence relatedness between PLE and ICP1. Notably, the 14-base pair sequence identical to the sequence in ICP1’s *pac* site is 100% conserved in the 10 known PLEs, and the 89-base pair sequence is 70-100% conserved between PLEs (Figure S1). PLEs’ sequence similarity to ICP1’s *pac* site is reminiscent of packaging sequence mimicry described in other satellites (4–6), suggesting PLE may use this sequence for PLE genome packaging, and therefore, it will be referred to as PLE’s putative *pac* site.

PLE’s putative *pac* site is 100% identical to ICP1’s across only 14 base pairs. Compared to other phage packaging mechanisms, 11-19 base pairs can be sufficient for TerS binding, but the surrounding sequence is often required for genome packaging (21–23). This could suggest that PLE mimics part of ICP1’s *pac* site to hijack ICP1’s TerS for packaging initiation or that PLE uses its own TerS that recognizes the similar but distinct sequence. Because ICP1’s TerS has not been identified, we could not look for predicted structural homologs in PLE. Instead, to identify candidates for PLE’s TerS, we used the observation that several phages encode their TerS near their *pac* site (24–26). The putative *pac* site in PLE lies between the origin of replication (*ori*) and 628 base pairs upstream of a four-gene cluster encoding three uncharacterized genes, *orf5, orf4, orf3,* and the characterized gene encoding the transcription factor CapR (18) (Figure 1B). During the late stages of ICP1 infection, when genome packaging occurs, this PLE gene cluster is expressed (27). Bioinformatic analysis of Orf3 (WP_002040293), Orf4 (WP_002040292), and Orf5 (WP_002040290) revealed no similarity with previously studied proteins (no E-values <1 by BLASTP or HHPRED). The fact that these genes are expressed when genome packaging occurs, that there is precedent for other phages to encode their *pac* sites within (24) or upstream (25, 26) of their *terS* gene, and the common clustering of genes with related functions within genomes led us to focus on these three uncharacterized genes as candidate PLE packaging proteins.

### PLE’s Orf3 inhibits ICP1’s TerL

Inhibitors of host phage genome packaging are a conserved feature of several satellite families (7–9). We hypothesized that one of the candidate PLE packaging proteins would inhibit ICP1 genome packaging and, therefore, inhibit plaque formation and that this would be a conserved feature found in all PLEs. Notably, all 10 known PLEs encode alleles of *orf3*, *orf4*, and *orf5* in the same syntenic organization (Figure 2A). Interestingly, there are variants of each gene that cluster into two groups based on amino acid similarity (Figure 2A and Figure S2). Starting with PLE1, we addressed the inhibitory effect of each candidate PLE packaging gene on ICP1 by individually expressing each gene on a plasmid during phage infection and quantifying the efficiency of plaquing (EOP) relative to the permissive empty vector control. While Orf4 and Orf5 had no negative effect on phage production, Orf3 reduced ICP1’s EOP by 1000-fold (Figure 2B). PLE10 encodes similar Orf4 and Orf5 variants relative to PLE1 (84% and 92% amino acid identity, respectively), but a variant of Orf3 with only 20% identity. As PLE10 has been shown to inhibit ICP1 and transduce at similar frequencies as PLE1 (28), we tested Orf3_PLE10_ (WP_001050346), Orf4_PLE10_ (WP_000073388), and Orf5_PLE10_ (WP_000095573) for ICP1 inhibition. Despite the sequence variation, Orf3 _PLE10_ decreased ICP1’s EOP to a similar level as Orf3 _PLE1_ (Figure 2B). Orf4_PLE10_ and Orf5_PLE10_ did not inhibit ICP1, as expected, due to their similarity to the PLE1 variants (Figure S3C). These results identify Orf3 as a novel and conserved inhibitory factor encoded by PLEs, making Orf3 the third factor that PLE encodes to antagonize ICP1, in addition to previously described NixI and TcaP (12, 13).

**Figure 2.**
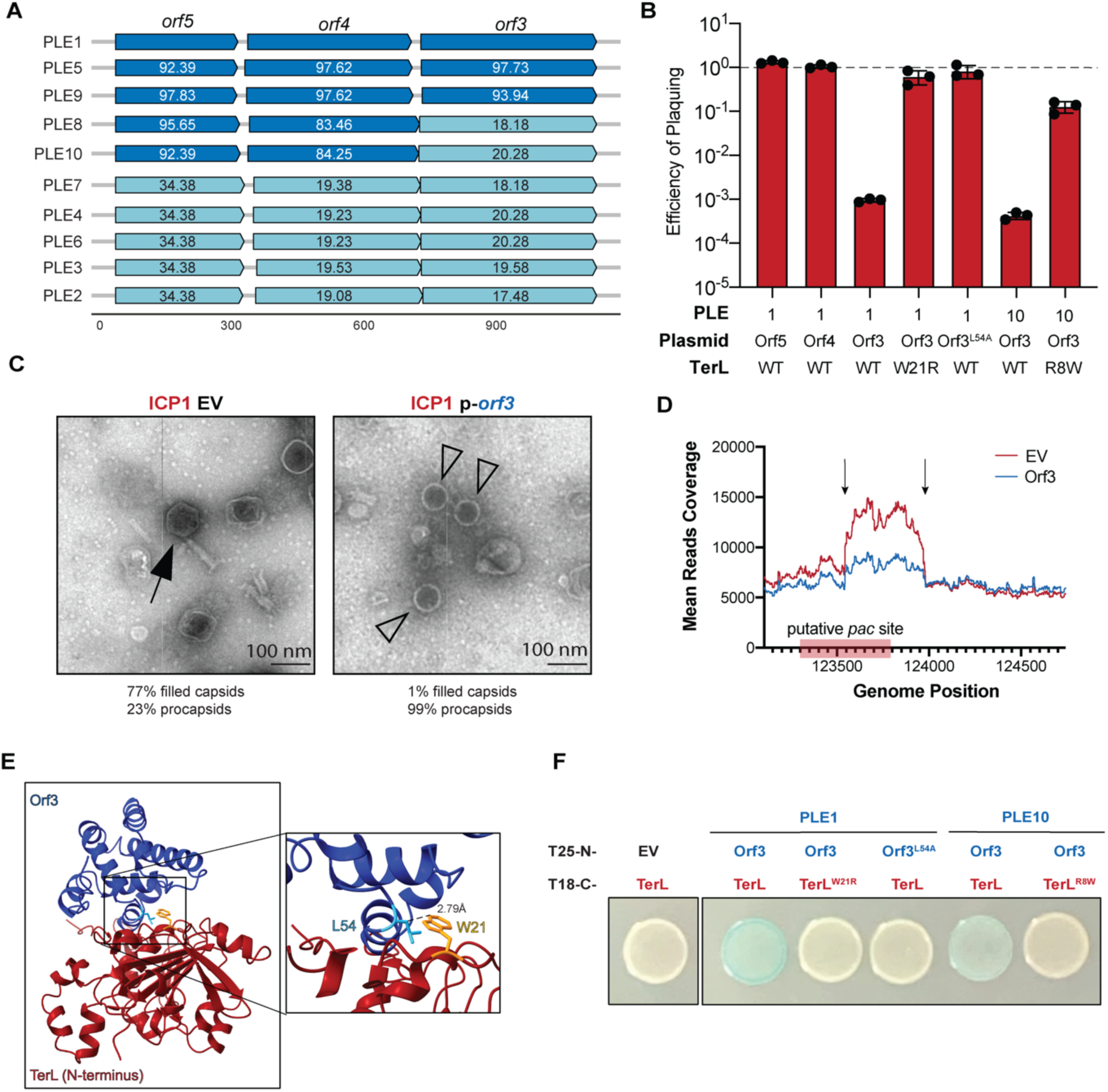
PLE-encoded Orf3 inhibits ICP1 genome packaging by directly binding to TerL. (A) Gene graphs of the putative packaging alleles from the 10 known PLEs. The scale indicated on the bottom is in base pairs. Numbers on the genes represent amino acid identity relative to PLE1 alleles. The two blues represent the two clusters of alleles, grouped by their amino acid similarity. (B) Efficiency of wild type (WT) or mutant ICP1 plaquing on *V. cholerae* strains expressing an empty vector, *orf5*, *orf4*, or *orf3* variants relative to an empty vector. The dashed line indicates an efficiency of 1 equal to the empty vector control. Each dot represents a biological replicate, and the error bars indicate the mean between the replicates (n=3). (C) Representative transmission electron micrographs of particles prepared on a *V. cholerae* strain expressing an empty vector or *orf3*. Arrows indicate mature ICP1 particles with encapsidated DNA and tails, while empty triangles indicate procapsids lacking DNA. Scale bar is 100 nm. Below: quantification of filled capsids and procapsids (EV: n=136, p-*orf3*: n=500). (D) Mean reads coverage (n=3) around ICP1’s *pac* site from DNA purified late during infection from strains expressing an empty vector (red) or Orf3_PLE1_ (blue). Arrows indicate cut sites. The red box indicates ICP1’s putative *pac* site. (E) AlphaFold2 predicted models of the N-terminus of TerL (red) and Orf3_PLE1_ (blue). Inset highlights the predicted interacting residues W21 (yellow) from TerL and L54 (cyan) from Orf3. The dashed line indicates the distance between the two indicated atoms. (F) Representative BACTH (bacterial adenylate cyclase two-hybrid) results from combinations of T25-N-Orf3 variants and T18-C-TerL variants (n=3). White indicates no interaction, while blue indicates an interaction between the two proteins.

To gain more insight into Orf3’s mechanism of inhibition, we assessed the morphology of particles produced from ICP1 infection in the presence of Orf3_PLE1_ by transmission electron microscopy (TEM). Virion assembly is a sequential process wherein procapsids and tails are produced in parallel while genomes are being replicated. Then genomes are packaged into the procapsids, resulting in capsid expansion, followed by addition of tails. Therefore, a disruption in one step of this process can result in a change of the abundance of an assembly precursor. In the Orf3-expressing strain, only 1% of capsids were filled with DNA, while the remaining 99% were procapsids, compared to 77% DNA-filled capsids and 23% procapsids produced from the empty vector control (Figure 2C). These data suggest there was a disruption after capsid assembly and before genome packaging, consistent with the hypothesis that Orf3 disrupts ICP1’s genome packaging. To assess this hypothesis further, we used a deep sequencing approach wherein we collected, purified, sequenced, and mapped total DNA from cultures at a late stage ICP1 of infection in the presence of Orf3 or an empty vector control. This approach allowed us to determine first if ICP1 genome replication was disrupted by Orf3 (which may explain the observed lack of DNA-filled particles) and second if genome packaging was occurring. First, a previous study showed that late during infection, ICP1 replicates by rolling circle replication, resulting in largely equal reads coverage across the genome (19). Concurrent with ICP1 replication, there is degradation of the *V. cholerae* chromosome, resulting in approximately 90% of the reads mapping to ICP1’s genome. Secondly, for headful packaging phages, when capsids are filled, there are usually two cleavage events that can be captured with this technique. Initially, the TerS-TerL-DNA complex is expected to cut near the *pac* site. Then TerL packages the DNA to fill the capsid, resulting in redundant packaging of ∼5-10% of the genome before TerL makes the second cut, which terminates packaging for that capsid. After cleavage, the TerL-DNA complex can then move to another procapsid and resume packaging, and this process can be repeated ∼10 times before TerL dissociates from the DNA (29). As such, the *pac* site and its surrounding sequence are more abundant than other sequences within capsids and are flanked by two cut sites. Between the Orf3-expressing and empty vector strains, reads coverage across ICP1’s genome was similar, indicating that Orf3 does not perturb ICP1 replication (Figure S3A&B). Most notably, however, reads coverage around the putative *pac* site was markedly depressed in the Orf3-expressing strain (Figure 2D and Figure S3A). In the Orf3-expressing strain, there was substantially lower coverage across a 577 base pair region between the cut sites and lower coverage across ∼500 base pairs upstream of the cut sites (Figure 2D). Collectively, these data are consistent with the hypothesis that Orf3 specifically inhibits ICP1 genome packaging.

Given that other satellites block genome packaging through inhibitors of their host phage’s TerS (7–9), we sought to identify Orf3’s target. To this end, we purified and sequenced the rare phage (n=1) that escaped inhibition by Orf3_PLE1_ and recovered the ability to plaque on this restrictive strain (Figure 2B and Table S3). Given the precedent in other satellite systems, we were surprised to find that this escape phage carried a single nonsynonymous mutation (W21R) in TerL (Table S3). To better understand how this suppressor substitution allowed for escape from Orf3_PLE1_, we used AlphaFold2 (30) to predict structures and potential interactions between TerL and Orf3_PLE1_ (Figure 2E). TerL’s structural prediction was of high confidence and revealed W21 in the N-terminal region, where an interaction with ICP1’s TerS complex would be expected (reviewed in 2). Although Orf3’s structural prediction was of very low confidence, the co-fold results predicted TerL’s W21 to interact with Orf3_PLE1_ at L54. This predicted interaction was supported by the lack of inhibition by Orf3_PLE1_^L54A^ (Figure 2B). Further, direct measurements of protein interactions between the Orf3_PLE1_ and TerL variants by bacterial adenylate cyclase two-hybrid (BACTH) assays showed the expected interactions: Orf3_PLE1_ and TerL^W21^ variants interacted, while TerL^W21R^ and Orf3_PLE1_^L54A^ variants did not interact with their wild type counterparts (Figure 2F). Next, we repeated these analyses for Orf3_PLE10_ which also restricted ICP1 plaquing (Figure 2B) but shares only 20% amino acid identity with Orf3_PLE1_ (Figure 2A). Despite the limited similarity in Orf3 proteins, phages (n=4) also escaped Orf3_PLE10_ with a single nonsynonymous mutation in TerL, however, substitutions were at a unique position (R8W) (Figure 2B, and Table S3). The TerL substitutions provided resistance only to the cognate Orf3 proteins on which they were selected (Figure S3D), consistent with direct interactions between these unique variants of Orf3 and TerL. We also detected interactions by BACTH between Orf3_PLE10_ and TerL (Figure S3E) and these findings are supported by a predicted interaction from AlphaFold2 co-fold analysis (Figure S3F). Although Orf4_PLE1_ and Orf5_PLE1_ did not inhibit ICP1 plaquing, given their proximity to the putative *pac* site, we tested for interactions with TerL, but no interactions were detected (Figure S3G). Interactions between the PLE-encoded putative packaging proteins were also not detected (Figure S3G). Taken together, these data indicate that PLE-encoded Orf3 targets and directly binds to ICP1’s TerL, and this interaction blocks ICP1 genome packaging.

### Orf3, Orf4, and Orf5 contribute to PLE transduction

Since PLE lacks its own TerL and is expected to be dependent ICP1’s TerL, and other phage satellites target their phage’s TerS (7–9), it was surprising that Orf3 targeted TerL. We reasoned that Orf3 could either be PLE’s TerS or function to specifically antagonize ICP1’s packaging, perhaps by blocking the interaction with ICP1’s TerS. To assess the role of Orf3 and the other candidate PLE packaging proteins in PLE’s genome packaging, we measured PLE transduction in the presence and absence of *orf3*, *orf4* and *orf5*. Given the more extensive characterization of PLE1 than PLE10, we used PLE1 for the remainder of this work and refer to this element as PLE for simplicity. Unlike a PLE(-) ICP1 infection, which results in the production of ICP1 progeny (Figure 3A), a PLE(+) infection produces PLE virions that can horizontally transfer the PLE genome to other cells (Figure 3B). Because PLE integrates following genome injection, a PLE-encoding an antibiotic resistance gene is used to select PLE(+) recipients in a transduction assay. In this manner, PLE transduction is quantified by the number of antibiotic-resistant recipient colonies following transduction. Relative to a WT PLE, PLEΔ*orf3* donors displayed a subtle 5-fold decrease in transduction efficiency (Figure 3C), suggesting Orf3 positively contributes to PLE transduction. The transduction of PLEΔ*orf4* and PLEΔ*orf5* revealed each has a more substantial 100-fold defect in transduction (Figure 3C), suggesting that these proteins also positively contribute to transduction. Plasmid-based expression of each gene complemented the transduction defects, ruling out polar effects within the operon (Figure 3D). To assess the interplay between these proteins, we constructed double and triple deletions and measured transduction. Interestingly, Orf3 was inhibitory to PLE transduction in genetic backgrounds without *orf4* and/or *orf5* (Figure 3C). For example, transduction of the *orf4* mutant improved nearly 20-fold when *orf3* was deleted. Additionally, when *orf3* was the only PLE packaging component expressed from the operon, transduction was just above the detection limit but improved over 120-fold for the triple deletion. These data rule against Orf3 as PLE’s TerS. Instead, they suggest that Orf3’s targeting of TerL can block PLE transduction and that Orf4 and Orf5 function additively to alleviate Orf3’s inhibitory effect on TerL to allow PLE to utilize this hijacked packaging motor. Further, these data demonstrate that despite the collective contribution to transduction, these three proteins are not wholly essential for transduction and suggest that PLE can also use a redundant, although less efficient, packaging strategy.

**Figure 3.**
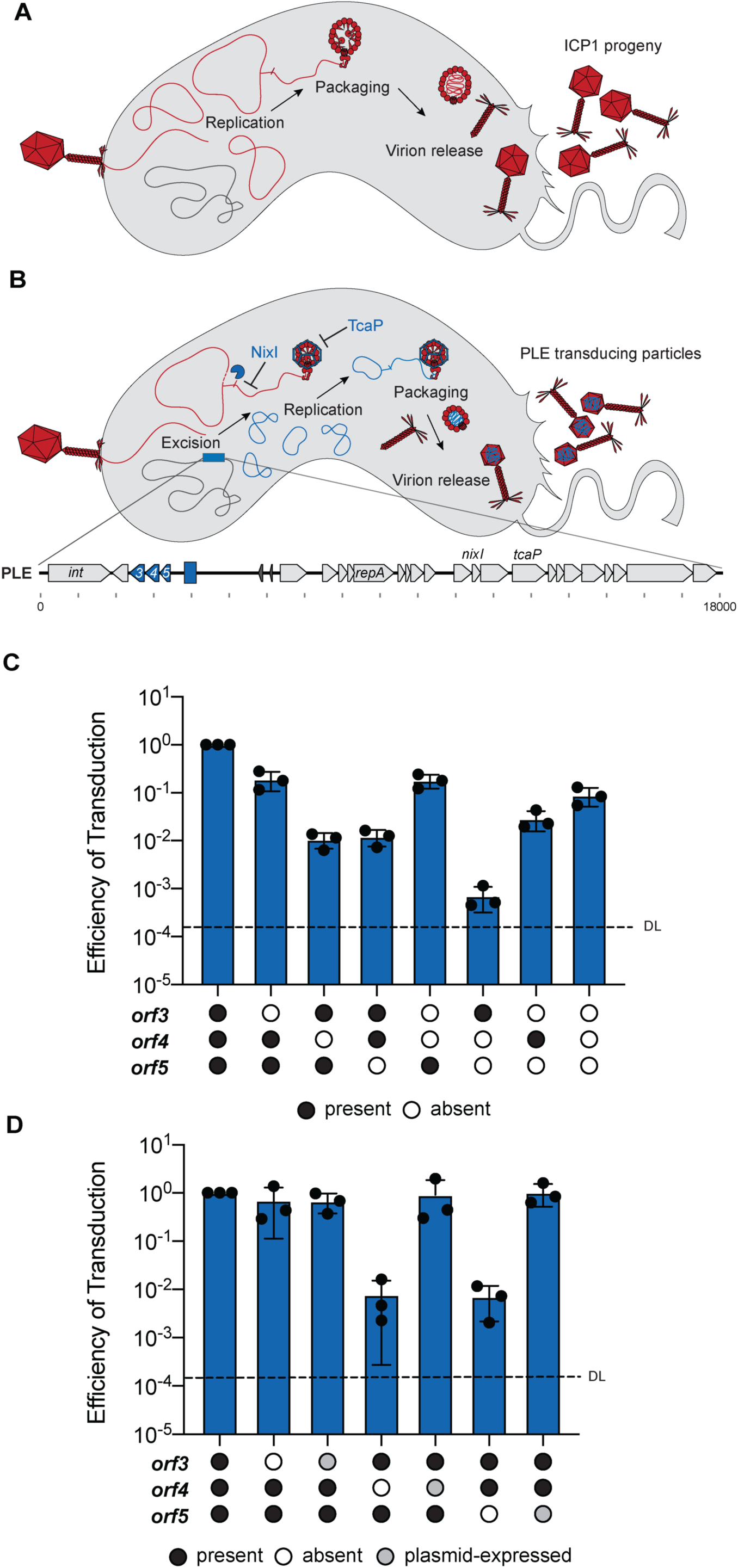
Orf4 and Orf5 overcome Orf3’s inhibition of PLE transduction. (A) A model of ICP1’s lifecycle, which results in ICP1 progeny. (B) A model of PLE’s lifecycle, highlighting excision, replication, packaging, and formation of transducing particles. The introduction of an antibiotic resistance gene after the last gene in PLE allows for the quantification of cells that acquire PLE from the transducing particles. PLE(+) infection does not produce ICP1 progeny. (C) Efficiency of transduction of PLE and PLE derivatives showing the effect of single, double, or triple deletions of *orf3*, *orf4*, and *orf5*, relative to the WT PLE. (D) Efficiency of transduction of PLE and PLE derivatives showing the complementation of single deletions of *orf3*, *orf4*, and *orf5*, relative to the WT PLE carrying an empty vector. Strains lacking a gene carry the empty vector. For (C) and (D) the dashed line indicates the detection limit and each dot represents a biological replicate, and the error bars indicate the mean between the replicates (n=3).

PLE’s parasitism of ICP1 is multifaceted, and many known and unknown factors can contribute to transduction. To eliminate factors unrelated to genome packaging specifically and more directly probe the roles of Orf3, Orf4, and Orf5 in genome packaging, we constructed a PLE that was <u>com</u>petent for <u>pac</u>kaging and transduction, named ComPacT-PLE (Figure 4A). ComPacT-PLE is a derivative of another reduced PLE element called midiPLE. midiPLE encodes attachment sites, the integrase, and an intergenic region with the putative *pac* site and the validated *ori*, and was shown to respond to ICP1 infection by excising and replicating, provided the PLE-encoded replication initiation factor RepA was expressed *in trans* (19). midiPLE, equivalent to ComPacT-PLEΔ*orf3*Δ*orf4*Δ*orf5*, does not transduce (Figure 4B). The addition of PLE’s putative packaging genes, *orf3, orf4,* and *orf5* to the element resulted in a 10,000-fold increase in transduction (Figure 4B). These data demonstrate that these are PLE’s packaging proteins which function to promote transduction. Further, single gene deletions in ComPacT-PLE resulted in 10- to 1,000-fold reductions in transduction efficiency (Figure 4B). The transduction defects could be rescued in each case by expression of the gene *in trans* (Figure 4B), demonstrating each protein contributes to transduction, as seen in PLE. Here, *orf3* and *orf5* mutants had more severe transduction defects than those observed in the context of PLE. ComPacT-PLE did not fully restrict ICP1 progeny production (Figure 4C) because this construct lacks PLE’s other inhibitory factors, notably NixI and TcaP. ICP1 was partially inhibited by ComPacT-PLEs that were Orf3 proficient (Figure 4C), in congruence with plasmid-based expression of Orf3 (Figure 2A). These data show that in its native context, Orf3 is inhibitory to ICP1.

**Figure 4.**
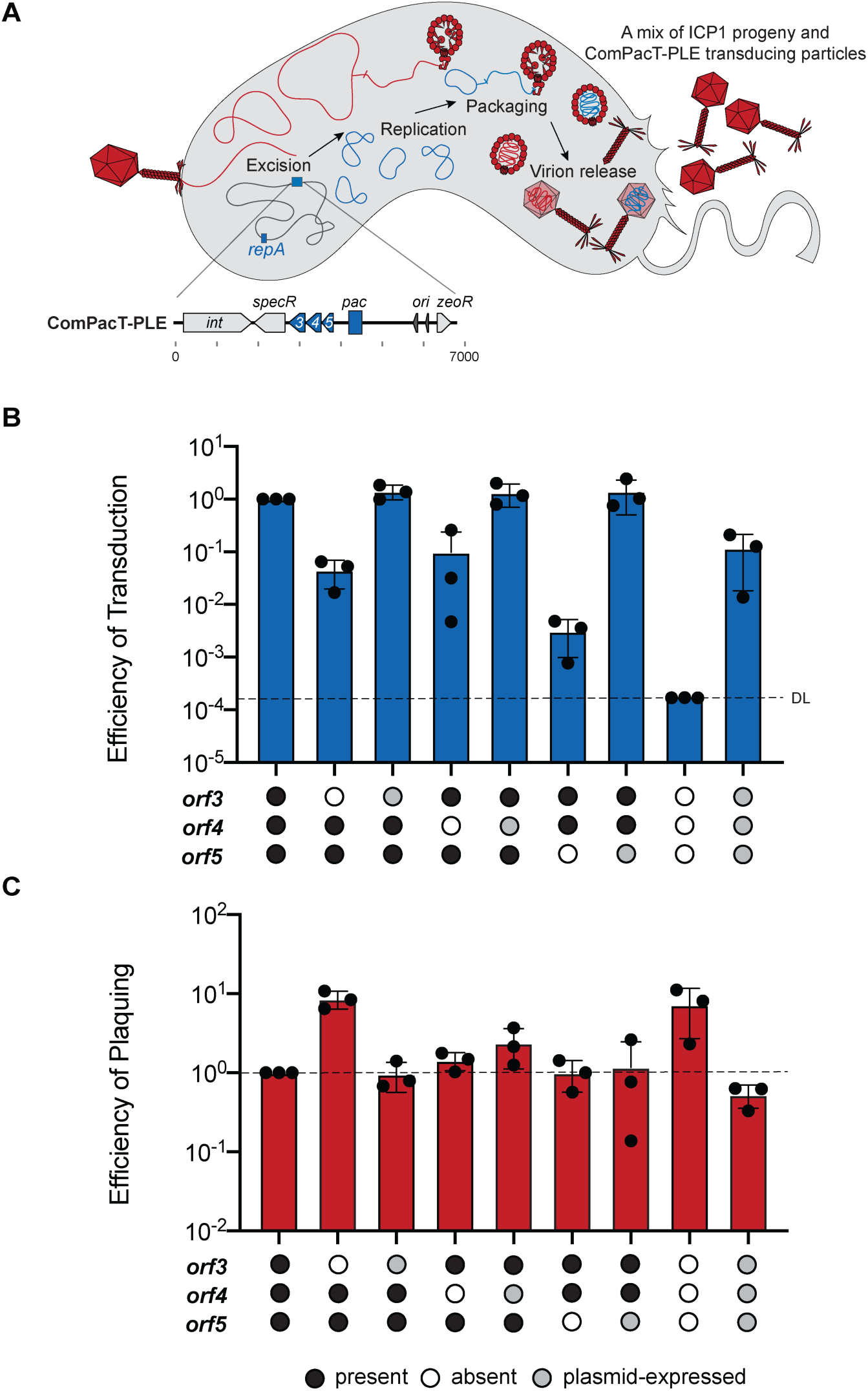
ComPacT-PLE uses Orf3, Orf4, and Orf5 to transduce. (A) A model showing ComPacT-PLE’s lifecycle, highlighting excision, replication, packaging, and virion release. The inset shows a gene map of ComPacT-PLE (drawn to scale). Note: *repA* is encoded in the *V. cholerae* chromosome to permit replication of ComPacT-PLE. (B) Efficiency of transduction of ComPacT-PLE and ComPacT-PLE derivatives showing the effect of single or triple deletions of *orf3*, *orf4*, and *orf5*, relative to WT ComPacT-PLE. The dashed line indicates the detection limit. Each dot represents a biological replicate, and the error bars indicate the mean between the replicates (n=3). Note that the strains lacking complementation carry an empty vector. (C) Efficiency of plaquing of the ICP1 produced from the same samples as in B, relative to WT ComPacT-PLE. Each dot represents a biological replicate, and the error bars indicate the mean between the replicates (n=3).

Paired with PLE transduction data (Figure 3), it is clear that Orf3 results in indiscriminate <u>g</u>enome <u>p</u>ackaging inhibition, so we have renamed it Gpi. Transduction data also show that Orf5 and Orf4 overcome Gpi’s inhibition to allow for <u>g</u>enome <u>p</u>ackaging <u>s</u>pecificity such that PLE genomes, but not ICP1 genomes, get packaged, and thus we renamed them GpsA and GpsB, respectively. The difference in the necessity for PLE’s packaging proteins between PLE and ComPacT-PLE is likely explained by the degree of competition between ICP1 and PLE because both entities rely on TerL. PLE decreases ICP1 genome replication with NixI, reducing the number of ICP1 genomes competing for TerL, an advantage not afforded to ComPacT-PLE. As such, ComPacT-PLE is more dependent on Gpi, GpsA, and GpsB than PLE is to capture TerLs from a limited pool and harness them to package satellite genomes.

### PLE requires the 14-base pairs that mimic ICP1’s *pac* site for transduction

Next, we set out to assess the contribution of PLE’s putative *pac* site to transduction using the minimal transducing unit, ComPacT-PLE. As 89 base pairs of PLE show similarity to a region of ICP1’s *pac* site (Figure 1C) and there is precedent for satellites to mimic their host phage’s packaging signals (4–6), we focused on this region. We constructed deletions in and around the putative *pac* site and measured the transduction of ComPacT-PLE. To facilitate the construction of the deletion strains of interest, an ampicillin resistance cassette was introduced downstream of the putative *pac* site, which only minimally decreased transduction, suggesting that the insertion site was outside of the *pac* site and that it did not significantly interfere with the neighboring *ori* or with packaging (Figure 5A). Next, we generated 89-50- and 14-base pair deletions (Δ89_1-89_, Δ50_39-89_, Δ14_75-89_, respectively) that included the 14-base pair sequence that is 100% identical to part of ICP1’s *pac* site (Figure 1C) and observed that these deletions abolished all transduction of ComPacT-PLE (Figure 5A). Deletions outside of the shared 14 base pairs resulted in detectable transduction. Transduction efficiency was least affected by the downstream deletion (Δ50_90-140_), while there was an approximately 10-fold reduction with the upstream deletion (Δ50_25-75_) within the rest of the putative *pac* site. These results demonstrate that the 14 base pairs shared with ICP1 and some of the upstream sequence contain PLE’s packaging signal and are required for transduction. To validate the role of PLE’s putative *pac* site in its native context, we also generated the 89- and 14-base pair deletions (Δ89_1-89_ and Δ14_75-89_, respectively) in PLE and measured transduction. Here, a spectinomycin resistance cassette was introduced in place of *capR* to facilitate the construction of the deletion strains. As previously reported (18), the deletion of *capR* had no impact on PLE transduction, while the deletions in the putative *pac* site resulted in ∼1000-fold reductions in transduction (Figure 5B). The 14-base pair deletion resulted in slightly higher transduction relative to the 89-base pair deletion, supporting that the sequence upstream of these shared base pairs contributes to PLE transduction. Similar to the transduction levels seen with deletions of PLE’s packaging proteins (Figure 3), PLE is more tolerant than ComPacT-PLE for partial deletions of its packaging signal (Figure 5), likely due to PLE’s superior inhibition of ICP1, which reduces competition for shared packaging resources.

**Figure 5.**
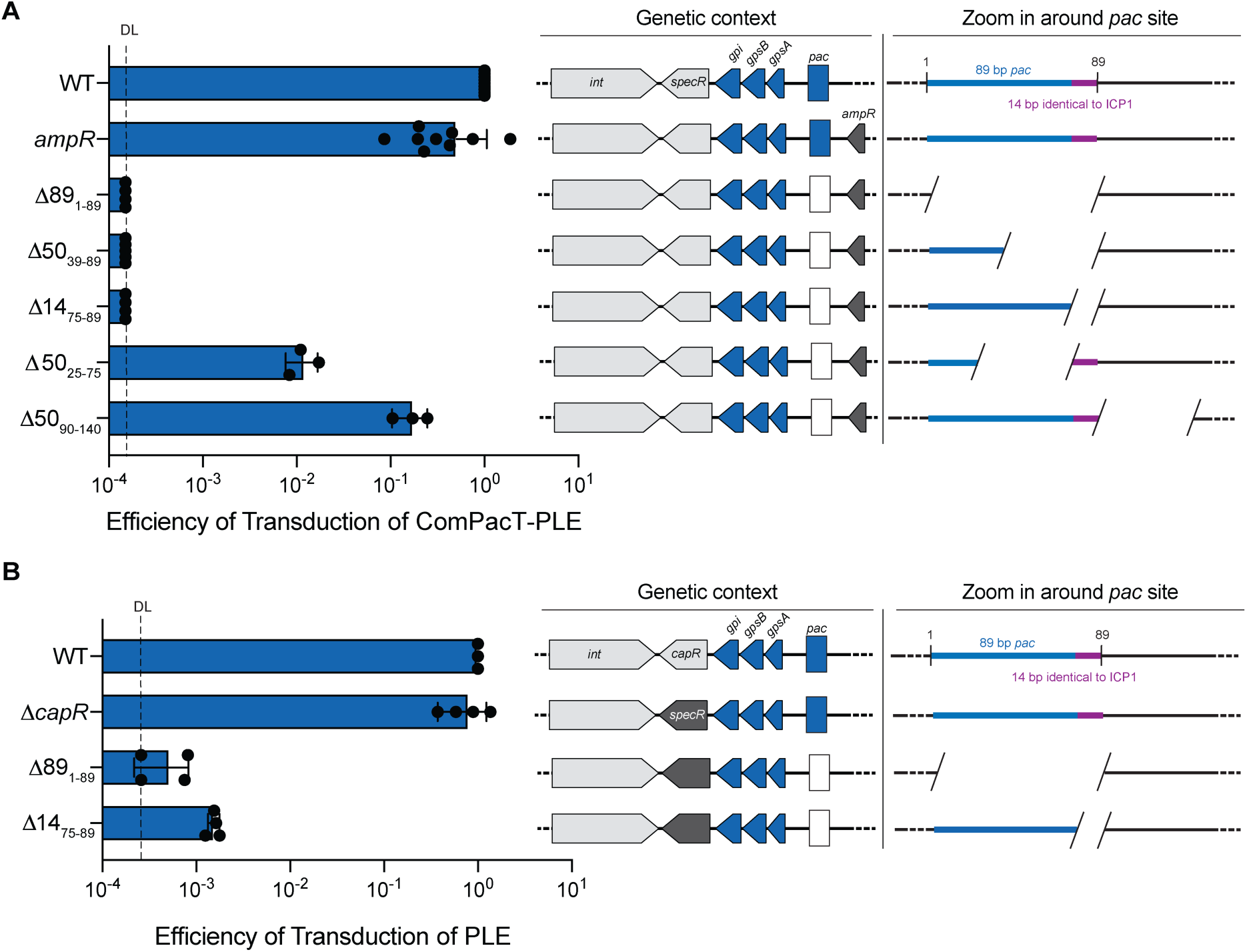
The 14 base pairs of PLE’s *pac* site that are shared with ICP1 are required for transduction. (A) Efficiency of transduction of ComPacT-PLE and its derivatives with deletions in the putative *pac* site. Deletions are labeled as positions relative to the 89-base pair putative *pac* site. Each dot represents a biological replicate, and the error bars indicate the mean between the replicates (n=3, 4, or 9, as indicated). The dashed line represents the detection limit. Diagrams to the right indicate 1) the genetic context of the 246-base pair region around the *pac* site (box) showing relevant antibiotic markers. An intact *pac* site is colored blue while deletions are indicated by a white box, and 2) a zoomed-in region around the relevant 89-base pair *pac* site is shown in blue and purple, where purple indicates the 14 base pairs that are identical to those in ICP1’s *pac* site (shown to scale). In both diagrams, the black line indicates the genome. For the zoom-in, the space between the hashes represents deletions. (B) Efficiency of transduction of PLE and its derivates with deletions in the putative *pac* site. Labels and diagrams are as in panel A.

## DISCUSSION

During an infection, phages and phage satellites need to differentiate their genomes to ensure specific packaging of their replicated genomes and exclude the host genome. Phage satellites that hijack their host phage’s capsids for transduction must compete for the limited resource of empty procapsids. Often, this requires manipulation of components of the packaging machinery, which are stolen from their host phages. Here, we report that the *V. cholerae* phage satellite of ICP1, PLE, uses a multifaceted strategy involving *pac* site mimicry and three proteins to coordinate the packaging of its genome. One of these proteins, Gpi, directly antagonizes genome packaging by binding to TerL. To our knowledge, this is the first example of a satellite directly inhibiting the large subunit rather than the small subunit of the host phage’s terminase complex. Through currently unknown mechanisms, GpsA and GpsB overcome the inhibitory function on TerL imposed by Gpi to specifically allow for PLE genome packaging and transduction. Surprisingly, we find that despite their sufficiency and necessity for transduction in ComPacT-PLE, the three packaging proteins are somewhat dispensable for transduction of an otherwise complete PLE. We anticipate that the additional anti-ICP1 factors encoded in the rest of PLE contribute to limiting ICP1 genomes and thus allow for PLE packaging. We favor a model (further described below and summarized in Figure 6) wherein PLE’s *pac* site mimicry contributes to packaging in these conditions.

**Figure 6.**
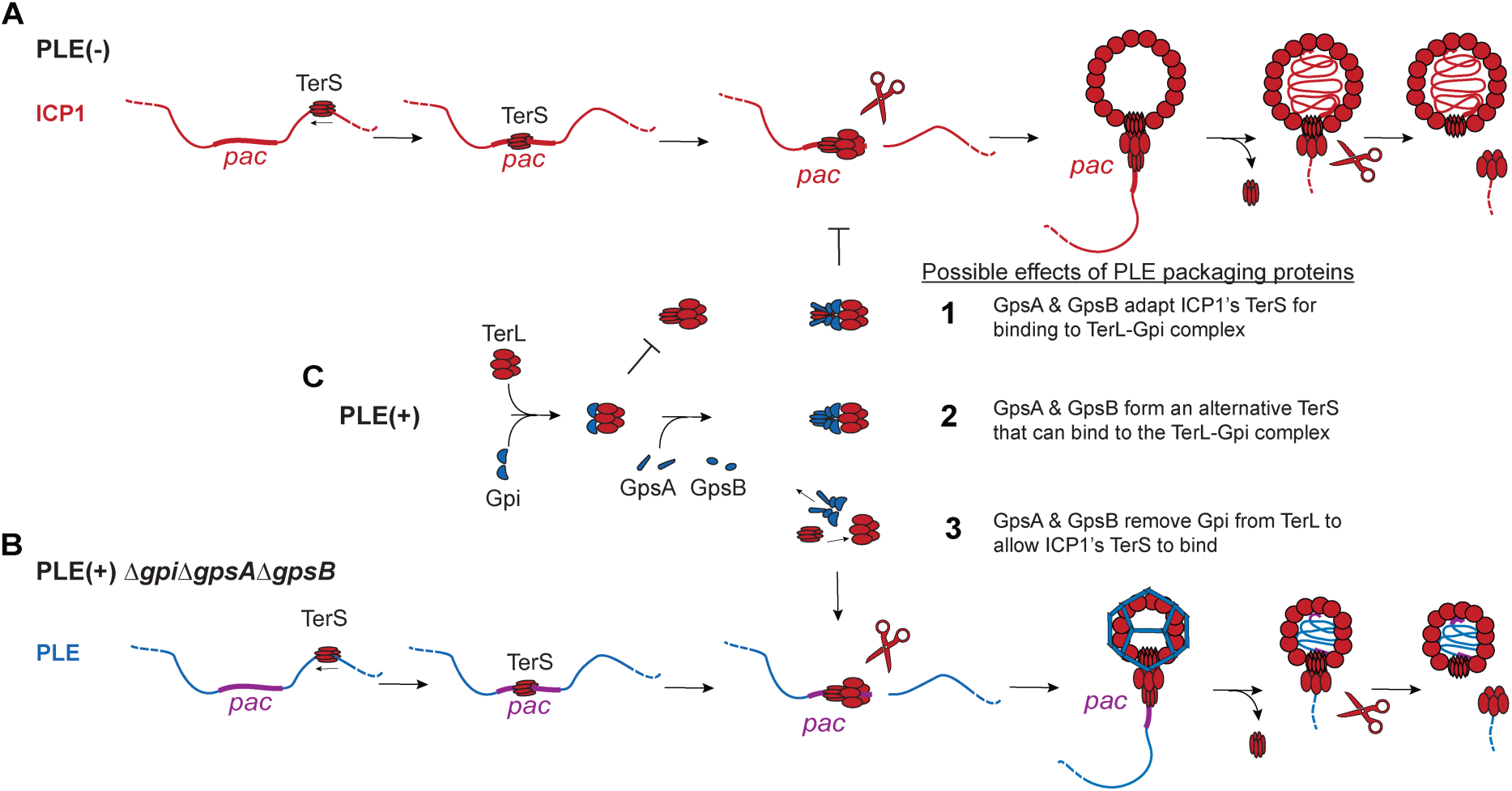
Possible models of ICP1 and PLE packaging. A) In a PLE(-) infection, ICP1’s TerS scans the concatemeric ICP1 DNA for the *pac* site. When bound to the *pac* site, TerS recruits TerL, which cleaves the genome, then transports the TerL-TerS-DNA complex to a procapsid. TerS presumably dissociates, and TerL translocates the DNA until the capsid is full. DNA is cleaved, and the TerL-DNA complex dissociates, primed to resume packaging into another empty procapsid. B) In the absence of PLE packaging proteins, PLE’s concatemeric DNA is similarly scanned by ICP1’s TerS for the sequence that mimics ICP1’s *pac* site (purple). PLE’s genome is packaged into TcaP-remodeled capsids in a similar manner described for ICP1’s packaging. C) In the presence of PLE’s packaging proteins, Gpi binds to TerL, presumably blocking an interaction between TerL and TerS. GpsA and GpsB may overcome Gpi’s inhibition of TerL through one of the following mechanisms: 1) by functioning as adapters to ICP1’s TerS such that this complex can bind to Gpi-TerL, 2) by functioning as an alternative TerS that has specificity for PLE’s *pac* site and is able to interact with Gpi-TerL, or 3) by removing Gpi from TerL such that ICP1’s TerS can bind. Presumably, the terminase complex from one of the possible scenarios depicted in panel C would replace the ICP1 terminase complex shown on PLE’s genome in panel B to mediate PLE genome packaging. DNA and proteins from ICP1 and PLE are shown in red and blue, respectively.

PLE’s packaging strategy represents a novel combination of previously described satellite packaging strategies with some notable differences. The three characterized mechanisms for satellite package redirection are exemplified by P4 and select Phage Inducible Chromosome Islands (PICIs). These characterized satellites are not related but have converged on manipulation of several stages of the phage lifecycle, including genome packaging, to permit horizontal spread of the satellites. First, P4, SaPIbov5, and PICI (EfCIV583) use a somewhat passive strategy wherein they mimic their host phage’s packaging signals such that the phage’s TerS and TerL recognize the satellite genomes as phage genomes and package them (4–6). Unlike P4 and SaPIbov5, PLEs’ *pac* site mimicry, while necessary (Figure 5), is only sufficient for transduction if PLE is equipped with other inhibitors of ICP1 (Figure 3C and Figure 4B). Second, some satellites of the PICI family (exemplified by EcCICFT073) repurpose their host phage’s TerS by changing its conformation such that it recognizes and, therefore, packages PICI but not host phage genomes (7). Because the identity of ICP1’s TerS is unknown, we cannot rule out that PLE also uses ICP1’s TerS and perhaps alters its specificity in favor of PLE genomes. In favor of this model, when PLE lacks its packaging proteins it suffers a transduction defect of only 10-fold (Figure 3C), suggesting PLE could use ICP1’s machinery in this context (Figure 6B). Additionally, ComPacT-PLE equipped with Gpi and either GpsA or GpsB allows for more transduction and is more inhibitory to ICP1 compared to a ComPacT-PLEΔ*gpi*Δ*gpsA*Δ*gpsB* (Figure 4B&C). These data fit a model wherein PLE can use ICP1’s TerS and provide specificity for PLE genome packaging (Figure 6C). In the third satellite-packaging example, a subset of satellites within the family of PICIs, the *Staphylococcus aureus* Pathogenicity Islands (SaPIs), achieve satellite packaging by encoding their own TerS that recognizes their genomes. To coordinate inhibition of phage genome packaging, these SaPIs deploy an inhibitor, Ppi, that binds to their host phage’s TerS, presumably rendering the phage’s TerS incompetent for phage genome packaging (8, 9). Like Ppi, PLE’s Gpi binds to a component of ICP1’s packaging machinery to interfere with phage packaging. However, Gpi binds to ICP1’s TerL (Figure 2) rather than TerS, and because PLE is dependent on ICP1’s TerL, Gpi can also interfere with PLE packaging (Figure 3C). PLE achieves immunity to Gpi via GpsA and GpsB, but the mechanism of immunity remains enigmatic. Perhaps GpsA and GpsB function together or separately as a PLE-specific TerS (Figure 6C). In line with this prediction, bioinformatic analysis hints that GpsB may have a DNA-binding motif, a feature expected of TerS proteins (31). Alternatively, GpsA and/or GpsB could modulate Gpi’s binding to TerL, selectively removing Gpi to allow ICP1’s TerS to package PLE genomes (Figure 6C). Such a mechanism would be analogous SaPI-encoded modifier, PitM, that fine-tunes PitA-mediated inhibition of a phage factor to favor satellite transmission (32). Moving forward, it will be important to define the precise mechanisms of Gpi, GpsA, and GpsB in TerL inhibition and genome packaging in the ICP1-PLE phage-satellite system. As newly discovered satellites are characterized, it will be interesting to compare their genome packaging strategies to see if there is conservation or innovation in diverse systems.

PLEs and SaPIs inhibit and redirect other processes in the phage lifecycle, which collectively favor satellite production over phage production, so they are not solely reliant on packaging inhibitors (Figure 3) (8). PLEs are notable as they are the only currently known satellites that fully restrict their host phage’s progeny production. Known PLE-encoded ICP1 inhibitors NixI, TcaP, and Gpi, discovered here, each target a different step in the ICP1 lifecycle (12, 13). As expected, individual PLE gene deletions are insufficient for ICP1 to escape PLE (14). In line with this redundancy, the suppressors from phages that afford escape from individual inhibitors have not been observed in our collection of natural ICP1 isolates to date (33). Instead of ICP1 relying on simultaneous mutational escape against every inhibitory PLE factor, ICP1 contends with PLE by encoding nuclease effectors (CRISPR-Cas, Odn, and/or Adi) that target and degrade the PLE genome (11, 34, 35). While CRISPR-Cas and Adi allow ICP1 to escape PLE’s inhibition enough to form phage progeny, PLE progeny production is not completely abolished in these conditions (35, 36). The remaining PLE genomes that have escaped ICP1’s targeting must directly compete with ICP1 genomes for the limited resources of empty procapsids and packaging machinery. Here, Gpi likely plays a significant role in blocking ICP1 genome packaging, and GpsA and GpsB are important for redirecting packaging motors for PLEs’ use. As seen by the difference in transduction efficiency between ComPacT-PLE and PLE lacking PLE packaging proteins, ComPacT-PLE is unable to outcompete ICP1 for these resources and does not get packaged nor transduced. PLE, on the other hand, outcompetes ICP1 in part through NixI-mediated reduction in ICP1 genome replication (13). NixI reduces ICP1 concatemer formation and thus reduces the number of ICP1 genomes, decreasing competition between ICP1 and PLE genomes for packaging machinery. In this manner, NixI and PLE’s packaging proteins act synergistically to ensure PLE transmission and exclusion of ICP1 genomes into virions.

Because of NixI’s activity, PLEs that are already outcompeting ICP1 for shared resources (empty procapsids and TerL complexes) can afford to rely on mimicry of ICP1’s *pac* site. While the *pac* site mimicry strategy may be sufficient for transduction, it is not as efficient as Gpi-, GpsA-, and GpsB-mediated transduction. The functional conservation of PLE’s packaging proteins supports the idea that these packaging proteins provide an advantage to PLE. Genetic variation between PLEs in the *gpi-gpsA-gpsB* locus, however, suggests there is selective pressure on this PLE packaging module. Gpi homologs show functional similarities but distinct interactions with TerL (Figure 2 and Figure S3). Interestingly, of the known PLE-encoded inhibitory proteins, only Gpi shows such high levels of variation, while NixI and TcaP alleles are more similar across the genetically distinct PLEs (28). The targets of NixI, TcaP, and Gpi are conserved in our collection of 67 clinical ICP1 isolates (33). Why Gpi varies more than the other inhibitors is unclear but may hint that some ICP1s encode countermeasures against Gpi. If PLE faced evasive strategies employed by ICP1 to escape Gpi, it would be beneficial for PLE to have a backup strategy to ensure its genome gets packaged. By mimicking ICP1’s *pac* site, PLE achieves packaging in the absence of functional Gpi, GpsA, and GpsB. In line with mimicry as a backup strategy in the face of ICP1 evolution, PLE’s *pac* site also shows variation. Specifically, the *pac* site of the more contemporary PLE10 is 22% more identical to ICP1’s *pac* site than any other PLE (Figure S1). Compared to PLE1, PLE10 shares 44 more base pairs in a continuous stretch and has 17 more identical base pairs over the 89 positions. Presumably, PLE10’s highly similar *pac* site would have a higher affinity for ICP1’s TerS than other PLEs and therefore PLE10 would be able to efficiently hijack ICP1’s TerS, even in the absence of PLE’s packaging proteins. Since mutations in TerS that would block PLE10’s *pac* site binding would also likely disrupt binding to ICP1’s *pac* site, PLE’s mimicry safeguards this strategy against ICP1’s mutational escape. Unfortunately, without knowing the identity of ICP1’s TerS, we cannot test these hypotheses. Overall, a response to ICP1 counter-defense may explain the variations of packaging proteins and the shift towards a more similar *pac* site in PLE10. ICP1 counter-defense to Gpi would also help explain why PLE would encode both *pac* site mimicry paired with packaging redirecting proteins, a combination not seen in other characterized satellite systems.

Phage satellites can provide population-wide immunity from virulent phages by limiting their host phage production during infection. In the last several years, many anti-phage defense systems have been found in bacterial genomes using systematic approaches (37–39). Although satellites are generally thought of as selfishly transmitting elements, it is interesting to compare their anti-phage mechanisms to those from other defense systems. Defense mechanisms against phage can be broken down into two main categories: 1) those that eliminate the phage prior to lysis (cell protective defense), and 2) those that limit phage spread by killing the host prior to virion assembly (abortive infection). In the first category, there are nucleic acid targeting systems that block phage genome replication (such as restriction-modification and CRISPR-Cas systems). Secondly, many defenses trigger cell death by recognizing structural proteins (40–47), but only a handful of mechanisms have been shown to specifically interfere with distinct steps of virion assembly. Two recently described defense systems provide protection by blocking tail assembly (48, 49). The remainder of virion assembly inhibitors identified to date are found in the realm of phage satellites. Satellites interfere with late gene expression (18, 32, 50), inhibit phage-sized capsid assembly (12, 51–53), and interfere with genome packaging (7–9). Interestingly, despite the notable structural dissimilarity of TerSs (31), other phage satellites target phage TerSs. In fact, this variable target can limit the satellites’ ability to interfere with other phages (8). Given the high conservation of TerL across highly divergent phages and even eukaryotic viruses, it is surprising that this protein is not the target of more phage satellites. The bacterial Avs defense system exploits the structural conservation of TerL and portal proteins as triggers to activate cell death in response to infection by diverse phages (54). Therapeutically, the conservation of TerL structures across diverse viruses, and their critical role in virus production make TerLs promising drug targets. Recently, P4 phage satellites were shown to carry broad-acting defense systems (55), suggesting satellites can give themselves a selective advantage by providing protection to their bacterial host from diverse infecting phages distinct from those they have evolved to parasitize. It will be interesting to see if any of the recently identified but not yet characterized defense systems target genome packaging by interfering with TerL, and if so, if there are mechanistic commonalities with Gpi. Because virion assembly strategies across diverse phages are conserved, PLE serves as a tool to discover novel and naturally occurring anti-phage mechanisms. These discoveries add to our collective understanding of viral lifecycles and have the potential to reveal strategies for viral interventions.

## DATA AVAILABILITY

The sequencing data generated for this work have been deposited in the Sequence Read Archive database under BioProject accession codes PRJNA1097610 and PRJNA1097611.

## AUTHOR CONTRIBUTIONS

Caroline M. Boyd: Conceptualization, Investigation, Methodology, Formal analysis, Visualization, Validation, Writing—original draft. Kimberley D. Seed: Conceptualization, Supervision, Writing—review & editing, Funding acquisition.

## ACKNOWLEDGEMENTS

We thank Seed Lab members, especially Kishen Patel, for their useful insights and feedback on this work. We also thank the helpful staff in the electron microscopy facility at the University of California, Berkeley.

## FUNDING

This publication was made possible by Grant Number R01AI127652 to K.D.S from the National Institute of Allergy and Infectious Diseases and the National Institute of Health NRSA Trainee Fellowship [5T32 CH132022 to C.M.B, in part.]. Its contents are solely the responsibility of the authors and do not necessarily represent the official views of the National Institute of Allergy and Infectious Diseases or National Institute of Health. K.D.S. holds an Investigators in the Pathogenesis of Infectious Disease Award from the Burroughs Wellcome Fund. Funding for open access charge: National Institute of Allergy and Infectious Diseases.

## CONFLICT OF INTEREST

The authors declare no conflicts of interest.

## Notes

### Competing Interest Statement

The authors have declared no competing interest.

## REFERENCES

1. de Sousa, J.A.M., Fillol-Salom, A., Penadés, J.R. and Rocha, E.P.C. (2023) Identification and characterization of thousands of bacteriophage satellites across bacteria. Nucleic Acids Research, 51, 2759–2777.

2. Catalano, C.E. and Morais, M.C. (2021) Viral genome packaging machines: Structure and enzymology. In The Enzymes. Elsevier, 50, 369–413.

3. Rao, V.B. and Feiss, M. (2015) Mechanisms of DNA Packaging by Large Double-Stranded DNA Viruses. Annual Review of Virology, 2, 351–378.

4. Martínez-Rubio, R., Quiles-Puchalt, N., Martí, M., Humphrey, S., Ram, G., Smyth, D., Chen, J., Novick, R.P. and Penadés, J.R. (2017) Phage-inducible islands in the Gram-positive cocci. ISME Journal, 11, 1029–1042.

5. Quiles-Puchalt, N., Carpena, N., Alonso, J.C., Novick, R.P., Marina, A. and Penadés, J.R. (2014) Staphylococcal pathogenicity island DNA packaging system involving cos-site packaging and phage-encoded HNH endonucleases. Proceedings of the National Academy of Sciences of the United States of America, 111, 6016–6021.

6. Ziermann, R. and Calendar, R. (1990) Characterization of the cos sites of bacteriophages P2 and P4. Gene, 96, 9–15.

7. Fillol-Salom, A., Bacarizo, J., Alqasmi, M., Ciges-Tomas, J.R., Martínez-Rubio, R., Roszak, A.W., Cogdell, R.J., Chen, J., Marina, A. and Penadés, J.R. (2019) Hijacking the Hijackers: Escherichia coli Pathogenicity Islands Redirect Helper Phage Packaging for Their Own Benefit. Molecular Cell, 75, 1020–1030.

8. Ram, G., Chen, J., Kumar, K., Ross, H.F., Ubeda, C., Damle, P.K., Lane, K.D., Penadfes, J.R., Christie, G.E. and Novick, R.P. (2012) Staphylococcal pathogenicity island interference with helper phage reproduction is a paradigm of molecular parasitism. Proceedings of the National Academy of Sciences of the United States of America, 109, 16300–16305.

9. Ubeda, C., Olivarez, N.P., Barry, P., Wang, H., Kong, X., Matthews, A., Tallent, S.M., Christie, G.E. and Novick, R.P. (2009) Specificity of staphylococcal phage and SaPI DNA packaging as revealed by integrase and terminase mutations. Molecular Microbiology, 72, 98–108.

10. O’Hara, B.J., Barth, Z.K., McKitterick, A.C. and Seed, K.D. (2017) A highly specific phage defense system is a conserved feature of the Vibrio cholerae mobilome. PLoS Genetics, 13, 1–17.

11. Seed, K.D., Lazinski, D.W., Calderwood, S.B. and Camilli, A. (2013) A bacteriophage encodes its own CRISPR/Cas adaptive response to evade host innate immunity. Nature, 494, 489–491.

12. Boyd, C.M., Subramanian, S., Dunham, D.T., Parent, K.N. and Seed, K.D. (2024) A Vibrio cholerae viral satellite maximizes its spread and inhibits phage by remodeling hijacked phage coat proteins into small capsids. eLife, 12, RP87611, 1–25.

13. LeGault, K.N., Barth, Z.K., DePaola, P. and Seed, K.D. (2022) A phage parasite deploys a nicking nuclease effector to inhibit viral host replication. Nucleic Acids Research, 1–17.

14. Hays, S.G. and Seed, K.D. (2020) Dominant Vibrio cholerae phage exhibits lysis inhibition sensitive to disruption by a defensive phage satellite. eLife, 9, 1–24.

15. Dalia, A.B., Lazinski, D.W. and Camilli, A. (2014) Identification of a Membrane-Bound Transcriptional Regulator That Links Chitin and Natural Competence in Vibrio cholerae. mBio, 5, 1–7.

16. Box, A.M., McGuffie, M.J., O’Hara, B.J. and Seed, K.D. (2016) Functional analysis of bacteriophage immunity through a Type I-E CRISPR-Cas system in Vibrio cholerae and its application in bacteriophage genome engineering. Journal of Bacteriology, 198, 578–590.

17. Ouellette, S.P., Karimova, G., Davi, M. and Ladant, D. (2017) Analysis of Membrane Protein Interactions with a Bacterial Adenylate Cyclase–Based Two-Hybrid (BACTH) Technique. CP Molecular Biology, 118, 20.12.1-20.12.24.

18. Netter, Z., Boyd, C.M., Silvas, T.V. and Seed, K.D. (2021) A phage satellite tunes inducing phage gene expression using a domesticated endonuclease to balance inhibition and virion hijacking. Nucleic Acids Research, 49, 1–16.

19. Barth, Z.K., Silvas, T.V., Angermeyer, A. and Seed, K.D. (2019) Genome replication dynamics of a bacteriophage and its satellite reveal strategies for parasitism and viral restriction. Nucleic acids research, 48, 249–263.

20. Barth, Z.K., Dunham, D.T. and Seed, K.D. (2023) Nuclease genes occupy boundaries of genetic exchange between bacteriophages. NAR Genomics and Bioinformatics, 5, 1–17.

21. Bento, J.C., Lane, K.D., Read, E.K., Cerca, N. and Christie, G.E. (2014) Sequence determinants for DNA packaging specificity in the S. aureus pathogenicity island SaPI1. Plasmid, 71, 8–15.

22. Chechik, M., Greive, S.J., Antson, A.A. and Jenkins, H.T. (2023) Structure of HK97 small terminase:DNA complex unveils a novel DNA binding mechanism by a circular protein. bioRxiv. doi: 10.1101/2023.07.17.549218, pre-print: not peer-reviewed

23. Chen, J., Ram, G., Penadés, J.R., Brown, S. and Novick, R.P. (2015) Pathogenicity Island-Directed Transfer of Unlinked Chromosomal Virulence Genes. Molecular Cell, 57, 138–149.

24. Oliveira, L., Tavares, P. and Alonso, J.C. (2013) Headful DNA packaging: Bacteriophage SPP1 as a model system. Virus Research, 173, 247–259.

25. MacHattie, L.A. and Gill, G.S. (1977) DNA maturation by the “headful” mode in bacteriophage T1. Journal of Molecular Biology, 110, 441–465.

26. Roberts, M.D., Martin, N.L. and Kropinski, A.M. (2004) The genome and proteome of coliphage T1. Virology, 318, 245–266.

27. Barth, Z.K., Netter, Z., Angermeyer, A., Bhardwaj, P. and Seed, K.D. (2020) A family of viral satellites manipulates invading virus gene expression and affects cholera toxin mobilization. mSystems, 5, 1– 18.

28. Angermeyer, A., Hays, S.G., Nguyen, M.H.T., Johura, F., Sultana, M., Alam, M. and Seed, K.D. (2022) Evolutionary Sweeps of Subviral Parasites and Their Phage Host Bring Unique Parasite Variants and Disappearance of a Phage CRISPR-Cas System. mBio, 13, 1–16.

29. Tavares, P., Lurz, R., Stiege, A., Rückert, B. and Trautner, T.A. (1996) Sequential Headful Packaging and Fate of the Cleaved DNA Ends in Bacteriophage SPP1. Journal of Molecular Biology, 264, 954– 967.

30. Mirdita, M., Schütze, K., Moriwaki, Y., Heo, L., Ovchinnikov, S. and Steinegger, M. (2022) ColabFold: making protein folding accessible to all. Nat Methods, 19, 679–682.

31. Lokareddy, R.K., Hou, C.-F.D., Li, F., Yang, R. and Cingolani, G. (2022) Viral Small Terminase: A Divergent Structural Framework for a Conserved Biological Function. Viruses, 14, 1–18.

32. Ram, G., Chen, J., Ross, H.F. and Novick, R.P. (2014) Precisely modulated pathogenicity island interference with late phage gene transcription. Proc. Natl. Acad. Sci. U.S.A., 111, 14536–14541.

33. Boyd, C.M., Angermeyer, A., Hays, S.G., Barth, Z.K., Patel, K.M. and Seed, K.D. (2021) Bacteriophage ICP1: A Persistent Predator of. Annual Reviews of Virology, 12, 1–20.

34. Barth, Z.K., Nguyen, M.H.T. and Seed, K.D. (2021) A chimeric nuclease substitutes a phage CRISPR-Cas system to provide sequence-specific immunity against subviral parasites. eLife, 1-21.

35. Nguyen, M.H.T., Netter, Z., Angermeyer, A. and Seed, K.D. (2022) A phage weaponizes a satellite recombinase to subvert viral restriction. Nucleic Acids Research, 50, 11138–11153.

36. McKitterick, A.C., LeGault, K.N., Angermeyer, A., Alam, M. and Seed, K.D. (2019) Competition between mobile genetic elements drives optimization of a phage-encoded CRISPR-Cas system: Insights from a natural arms race. Philosophical Transactions of the Royal Society B: Biological Sciences, 374, 1–12.

37. Tesson, F., Hervé, A., Mordret, E., Touchon, M., d’Humières, C., Cury, J. and Bernheim, A. (2022) Systematic and quantitative view of the antiviral arsenal of prokaryotes. Nat Commun, 13, 1–10.

38. Payne, L.J., Todeschini, T.C., Wu, Y., Perry, B.J., Ronson, C.W., Fineran, P.C., Nobrega, F.L. and Jackson, S.A. (2021) Identification and classification of antiviral defence systems in bacteria and archaea with PADLOC reveals new system types. Nucleic Acids Research, 49, 10868–10878.

39. Doron, S., Melamed, S., Ofir, G., Leavitt, A., Lopatina, A., Keren, M., Amitai, G. and Sorek, R. (2018) Systematic discovery of antiphage defense systems in the microbial pangenome. Science, 359, 1–12.

40. Huiting, E., Cao, X., Ren, J., Athukoralage, J.S., Luo, Z., Silas, S., An, N., Carion, H., Zhou, Y., Fraser, J.S., et al. (2023) Bacteriophages inhibit and evade cGAS-like immune function in bacteria. Cell, 186, 864–876.

41. Stokar-Avihail, A., Fedorenko, T., Hör, J., Garb, J., Leavitt, A., Millman, A., Shulman, G., Wojtania, N., Melamed, S., Amitai, G., et al. (2023) Discovery of phage determinants that confer sensitivity to bacterial immune systems. Cell, 186, 1863–1876.

42. Garb, J., Lopatina, A., Bernheim, A., Zaremba, M., Siksnys, V., Melamed, S., Leavitt, A., Millman, A., Amitai, G. and Sorek, R. (2022) Multiple phage resistance systems inhibit infection via SIR2-dependent NAD+ depletion. Nat Microbiol, 7, 1849–1856.

43. Zhang, T., Tamman, H., Coppieters ’T Wallant, K., Kurata, T., LeRoux, M., Srikant, S., Brodiazhenko, T., Cepauskas, A., Talavera, A., Martens, C., et al. (2022) Direct activation of a bacterial innate immune system by a viral capsid protein. Nature, 612, 132–140.

44. Tal, N., Morehouse, B.R., Millman, A., Stokar-Avihail, A., Avraham, C., Fedorenko, T., Yirmiya, E., Herbst, E., Brandis, A., Mehlman, T., et al. (2021) Cyclic CMP and cyclic UMP mediate bacterial immunity against phages. Cell, 184, 5728–5739.

45. Schmitt, C.K. and Molineux, I.J. (1991) Expression of gene 1.2 and gene 10 of bacteriophage T7 is lethal to F plasmid-containing Escherichia coli. J Bacteriol, 173, 1536–1543.

46. Molineux, I.J., Schmitt, C.K. and Condreay, J.P. (1989) Mutants of bacteriophage T7 that escape F restriction. Journal of Molecular Biology, 207, 563–574.

47. Champness, W.C. and Snyder, L. (1982) The gol site: a cis-acting bacteriophage T4 Regulatory Region that can affect expression of all the T4 late genes. Journal of Molecular Biology, 155, 395– 407.

48. Patel, P.H., Taylor, V.L., Zhang, C., Getz, L.J., Fitzpatrick, A.D., Davidson, A.R. and Maxwell, K.L. (2024) Anti-phage defence through inhibition of virion assembly. Nat Commun, 15, 1–11.

49. Hör, J., Wolf, S.G. and Sorek, R. (2023) Bacteria conjugate ubiquitin-like proteins to interfere with phage assembly Microbiology, bioRxiv, doi: 10.1101/2023.09.04.556158, pre-print: not peer-reviewed.

50. Lindqvist, B.H., Deho, G. and Calendar, R. (1993) Mechanisms of genome propagation and helper exploitation by satellite phage P4. Microbiological Reviews, 57, 683–702.

51. Carpena, N., Manning, K.A., Dokland, T., Marina, A. and Penadés, J.R. (2016) Convergent evolution of pathogenicity islands in helper cos phage interference. Philosophical Transactions of the Royal Society B: Biological Sciences, 371, 1–9.

52. Damle, P.K., Wall, E.A., Spilman, M.S., Dearborn, A.D., Ram, G., Novick, R.P., Dokland, T. and Christie, G.E. (2012) The roles of SaPI1 proteins gp7 (CpmA) and gp6 (CpmB) in capsid size determination and helper phage interference. Virology, 432, 277–282.

53. Shore, D., Deho, G., Tsipis, J. and Goldstein, R. (1978) Determination of capsid size by satellite bacteriophage P4. Proceedings of the National Academy of Sciences of the United States of America, 75, 400–404.

54. Gao, L.A., Wilkinson, M.E., Strecker, J., Makarova, K.S., Macrae, R.K., Koonin, E.V. and Zhang, F. (2022) Prokaryotic innate immunity through pattern recognition of conserved viral proteins. *Science (New York*, N.Y.), 377, 1–15.

55. Rousset, F., Depardieu, F., Miele, S., Dowding, J., Laval, A.-L., Lieberman, E., Garry, D., Rocha, E.P.C., Bernheim, A. and Bikard, D. (2022) Phages and their satellites encode hotspots of antiviral systems. Cell Host & Microbe, 30, 740–753.

