## Supplementary material for "A phage satellite manipulates the viral DNA packaging motor to inhibit phage and promote satellite spread": Tables S1-S3, Figures S1-S3

**Supplementary Table S1: Strains used in this study**

| Strain | Description | Source |
|---|---|---|
| KDS6 | *V. cholerae* O1, El Tor biotype | Lab collection |
| KDS103 | *V. cholerae* E7946 containing PLE1::kanR[1] | (1) |
| KDS150 | *V. cholerae* E7946 PLE1::kanR Δ*orf3*::frt | This study |
| KDS151 | *V. cholerae* E7946 PLE1::kanR Δ*orf4*::frt | This study |
| CMB618 | *V. cholerae* E7946 PLE1::kanR Δ*orf5*::frt | This study |
| CMB656 | *V. cholerae* E7946 PLE1::kanR Δ*orf5-orf4*::frt | This study |
| CMB663 | *V. cholerae* E7946 PLE1::kanR Δ*orf4-orf3*::frt | This study |
| CMB620 | *V. cholerae* E7946 PLE1::kanR Δ*orf5-orf3*::frt | This study |
| CMB962 | *V. cholerae* E7946 PLE1::kanR Δ*orf5::frt*Δ*orf3* | This study |
| CMB578 | *V. cholerae* E7946 containing PLE1::kanR pKL06.2 $P_{tac}$-RiboE-empty | This study |
| CMB497 | *V. cholerae* E7946 PLE1::kanR Δ*orf3*::frt pKL06.2 $P_{tac}$-RiboE-*orf3* | This study |
| CMB896 | *V. cholerae* E7946 PLE1::kanR Δ*orf4*::frt pKL06.2 $P_{tac}$-RiboE-*orf4* | This study |
| CMB900 | *V. cholerae* E7946 PLE1::kanR Δ*orf5*::frt pKL06.2 $P_{tac}$-RiboE-*orf5* | This study |
| CMB496 | *V. cholerae* E7946 PLE1::kanR Δ*orf3*::frt pKL06.2 $P_{tac}$-RiboE-*empty* | This study |
| CMB895 | *V. cholerae* E7946 PLE1::kanR Δ*orf4*::frt pKL06.2 $P_{tac}$-RiboE- *empty* | This study |
| CMB899 | *V. cholerae* E7946 PLE1::kanR Δ*orf5*::frt pKL06.2 $P_{tac}$-RiboE- *empty* | This study |
| CMB3 | *V. cholerae* E7946 pKL06.2 $P_{tac}$-RiboE-empty | This study |
| KDS199 | *V. cholerae* E7946 pKL06.2 $P_{tac}$-RiboE-$orf3_{PLE1}$ | This study |
| CMB870/871 | *V. cholerae* E7946 pKL06.2 $P_{tac}$-RiboE-$orf3_{PLE1}^{L43A}$ | This study |
| KS2565 | *V. cholerae* E7946 pKL06.2 $P_{tac}$-RiboE-$orf3_{PLE10}$ | This study |
| KS1912 | *V. cholerae* E7946 pKL06.2 $P_{tac}$-RiboE-$orf4_{PLE1}$ | This study |
| KS1807 | *V. cholerae* E7946 pKL06.2 $P_{tac}$-RiboE-$orf5_{PLE1}$ | This study |
| CMB856/857 | *V. cholerae* E7946 pKL06.2 $P_{tac}$-RiboE-$orf4_{PLE10}$ | This study |
| CMB866/867 | *V. cholerae* E7946 pKL06.2 $P_{tac}$-RiboE-$orf5_{PLE10}$ | This study |
| CMB1018 | *V. cholerae* E7946 PLE1::kanR Δ*orf2*::specR | This study |
| CMB1020 | *V. cholerae* E7946 PLE1::kanR Δ*orf2*::specR Δ$pac_{89}$ | This study |
| CMB1027 | *V. cholerae* E7946 PLE1::kanR Δ*orf2*::specR Δ$pac_{14}$ | This study |
| CMB862 | *V. cholerae* E7946 *ΔlacZ*::*trimR* | This study |
| KDS2421 | *V. cholerae* E7946 *ΔlacZ*::*specR* | This study |
| CMB650 | *V. cholerae* E7946 ComPacT-PLE, *ΔlacZ*::$P_{tac}$-RiboE-*repA*, KanR, SpecR | This study |
| CMB672 | *V. cholerae* E7946 ComPacT-PLE, *ΔlacZ*::$P_{tac}$-RiboE-*repA*, pKL06.2 $P_{tac}$-RiboE-empty KanR, SpecR, CmR | This study |
| CMB676 | *V. cholerae* E7946 ComPacT-PLEΔ*orf3*::frt, *ΔlacZ*::$P_{tac}$-RiboE-*repA*, pKL06.2 $P_{tac}$-RiboE-*empty* KanR, SpecR, CmR | This study |

| Strain | Description | Source |
|---|---|---|
| CMB710 | *V. cholerae* E7946 ComPacT-PLEΔ*orf4*::frt, Δ*lacZ*::$P_{tac}$-RiboE-*repA*, pKL06.2 $P_{tac}$-RiboE-*empty* KanR, SpecR, CmR | This study |
| CMB714 | *V. cholerae* E7946 ComPacT-PLEΔ*orf5*::frt, Δ*lacZ*::$P_{tac}$-RiboE-*repA*, pKL06.2 $P_{tac}$-RiboE-*empty* KanR, SpecR, CmR | This study |
| CMB684 | *V. cholerae* E7946 ComPacT-PLEΔ*orf3-orf5*::frt, Δ*lacZ*::$P_{tac}$-RiboE-*repA*, pKL06.2 $P_{tac}$-RiboE-*empty* KanR, SpecR, CmR | This study |
| CMB678 | *V. cholerae* E7946 ComPacT-PLEΔ*orf3*::frt, Δ*lacZ*::$P_{tac}$-RiboE-*repA*, pKL06.2 $P_{tac}$-RiboE-*orf3* KanR, SpecR, CmR | This study |
| CMB712 | *V. cholerae* E7946 ComPacT-PLEΔ*orf4*::frt, Δ*lacZ*::$P_{tac}$-RiboE-*repA*, pKL06.2 $P_{tac}$-RiboE-*orf4* KanR, SpecR, CmR | This study |
| CMB716 | *V. cholerae* E7946 ComPacT-PLEΔ*orf5*::frt, Δ*lacZ*::$P_{tac}$-RiboE-*repA*, pKL06.2 $P_{tac}$-RiboE-*orf5* KanR, SpecR, CmR | This study |
| CMB688 | *V. cholerae* E7946 ComPacT-PLEΔ*orf3-orf5*::frt, Δ*lacZ*::$P_{tac}$-RiboE-*repA*, pKL06.2 $P_{tac}$-RiboE-*orf5-orf4-orf3* KanR, SpecR, CmR | This study |
| CMB703 | *V. cholerae* E7946 ComPacT-PLE *pac*::ampR, Δ*lacZ*::$P_{tac}$-RiboE-*repA*, KanR, SpecR, AmpR | This study |
| CMB837 | *V. cholerae* E7946 ComPacT-PLE Δ89$pac_{1-89}$, Δ*lacZ*::$P_{tac}$-RiboE-*repA*, KanR, SpecR, AmpR | This study |
| CMB839 | *V. cholerae* E7946 ComPacT-PLE Δ50$pac_{74-89}$, Δ*lacZ*::$P_{tac}$-RiboE-*repA*, KanR, SpecR, AmpR | This study |
| CMB880 | *V. cholerae* E7946 ComPacT-PLE Δ$pac_{14}$, Δ*lacZ*::$P_{tac}$-RiboE-*repA*, KanR, SpecR, AmpR | This study |
| CMB887 | *V. cholerae* E7946 ComPacT-PLE Δ50$pac_{25-75}$, Δ*lacZ*::$P_{tac}$-RiboE-*repA*, KanR, SpecR, AmpR | This study |
| CMB889 | *V. cholerae* E7946 ComPacT-PLE Δ50$pac_{90-140}$, Δ*lacZ*::$P_{tac}$-RiboE-*repA*, KanR, SpecR, AmpR | This study |
| CMB465 | *E. coli* XL1-Blue T-25-N-linker-$orf3_{PLE1}$ | This study |
| CMB457 | *E. coli* XL1-Blue T-25-C-linker-$orf3_{PLE1}$ | This study |
| CMB461 | *E. coli* XL1-Blue T-18-N-linker-$orf3_{PLE1}$ | This study |
| CMB453 | *E. coli* XL1-Blue T-18-C-linker-$orf3_{PLE1}$ | This study |
| CMB470 | *E. coli* XL1-Blue T-25-N-linker-$orf3_{PLE10}$ | This study |
| CMB468 | *E. coli* XL1-Blue T-25-C-linker-$orf3_{PLE10}$ | This study |
| CMB469 | *E. coli* XL1-Blue T-18-N-linker-$orf3_{PLE10}$ | This study |
| CMB467 | *E. coli* XL1-Blue T-18-C-linker-$orf3_{PLE10}$ | This study |
| CMB881 | *E. coli* XL1-Blue T-25-N-linker-$orf3_{PLE1}{}^{L54A}$ | This study |
| CMB882 | *E. coli* XL1-Blue T-25-C-linker-$orf3_{PLE1}{}^{L54A}$ | This study |
| CMB883 | *E. coli* XL1-Blue T-18-N-linker-$orf3_{PLE1}{}^{L54A}$ | This study |
| CMB884 | *E. coli* XL1-Blue T-18-C-linker-$orf3_{PLE1}{}^{L54A}$ | This study |
| CMB459 | *E. coli* XL1-Blue T-25-N-linker-*terL* | This study |

| Strain | Description | Source |
|---|---|---|
| CMB501 | *E. coli* XL1-Blue T-25-N-linker-$terL^{W21R}$ | This study |
| CMB502 | *E. coli* XL1-Blue T-25-N-linker-$terL^{R8W}$ | This study |
| CMB451 | *E. coli* XL1-Blue T-18-C-linker-*terL* | This study |
| CMB480 | *E. coli* XL1-Blue T-18-C-linker-$terL^{W21R}$ | This study |
| CMB479 | *E. coli* XL1-Blue T-18-C-linker-$terL^{R8W}$ | This study |
| CMB622 | *E. coli* XL1-Blue T-25-N-linker-$orf4_{PLE1}$ | This study |
| CMB632 | *E. coli* XL1-Blue T-25-C-linker-$orf4_{PLE1}$ | This study |
| CMB627 | *E. coli* XL1-Blue T-18-N-linker-$orf4_{PLE1}$ | This study |
| CMB637 | *E. coli* XL1-Blue T-18-C-linker-$orf4_{PLE1}$ | This study |
| CMB466 | *E. coli* XL1-Blue T-25-N-linker-$orf5_{PLE1}$ | This study |
| CMB458 | *E. coli* XL1-Blue T-25-C-linker-$orf5_{PLE1}$ | This study |
| CMB462 | *E. coli* XL1-Blue T-18-N-linker-$orf5_{PLE1}$ | This study |
| CMB454 | *E. coli* XL1-Blue T-18-C-linker-$orf5_{PLE1}$ | This study |
| ICP1 | ICP1_2018_Mat_B accession: MW794183 | (2) |
| KSphi151 | ICP1_2018_Mat_B $TerL^{W21R}$ | This study |
| KSphi152 | ICP1_2018_Mat_B $TerL^{R8W}$ | This study |
| CMBphi34 | ICP1_2018_Mat_B ΔCRISPR ΔCas2-3 | This study |
| CMBphi35 | ICP1_2018_Mat_B ΔCRISPR ΔCas2-3 $TerL^{W21R}$ | This study |
| CMBphi36 | ICP1_2018_Mat_B $TerL^{R8W,W21R}$ | This study |

[1]KanR = Kanamycin resistance cassette, SpecR = Spectinomycin resistance cassette, TrimR = Trimethoprim resistance cassette, AmpR = Ampicillin resistance cassette

**Supplementary Table S2. Top five BLASTN hits for PLE1 against ICP1**

| Query | Hit | ID | Description | E-value | HSP Start | HSP end | HSP length | Query start | Query end | % identity | %positive |
|---|---|---|---|---|---|---|---|---|---|---|---|
| **PLE1** | ICP1 | *capR* | gp165 | 8.75E-05 | 91528 | 91388 | 141 | 2176 | 2316 | 68.75 | 68.75 |
| **PLE1** | ICP1 | put. *pac* | pac site | 0.045334 | 399 | 486 | 88 | 3975 | 4063 | 69.66 | 69.66 |
| **PLE1** | ICP1 | NA[1] | NA | 0.158232 | 87755 | 87720 | 36 | 14855 | 14890 | 80.56 | 80.56 |
| **PLE1** | ICP1 | NA | NA | 0.158232 | 12513 | 12516 | 33 | 12612 | 12644 | 81.82 | 81.82 |
| **PLE1** | ICP1 | NA | NA | 0.552285 | 79488 | 79512 | 25 | 2128 | 2152 | 88 | 88 |

[1]NA= not applicable

**Supplementary Table S3: Escape phage results**

| Phage[1] | Escape from | Position | SNP[1] | Gene | Protein |
|---|---|---|---|---|---|
| KSphi151 | $Orf3_{PLE1}$ | 79,700 | A → G | *gp129* | $TerL^{W21R}$ |
| KSphi152 | $Orf3_{PLE10}$ | 79,809 | T → A | *gp129* | $TerL^{R8W}$ |
| KSphi153 | $Orf3_{PLE10}$ | 79,809 | T → A | *gp129* | $TerL^{R8W}$ |
| KSphi154 | $Orf3_{PLE10}$ | 79,809 | T → A | *gp129* | $TerL^{R8W}$ |
| KSphi155 | $Orf3_{PLE10}$ | 79,809 | T → A | *gp129* | $TerL^{R8W}$ |

[1]These phages are derivatives of ICP1_2018_Mat_B accession MW794183
[2]All single nucleotide polymorphisms (SNPs) at 100% frequency in the sequenced population are shown.

### Supplementary Figures

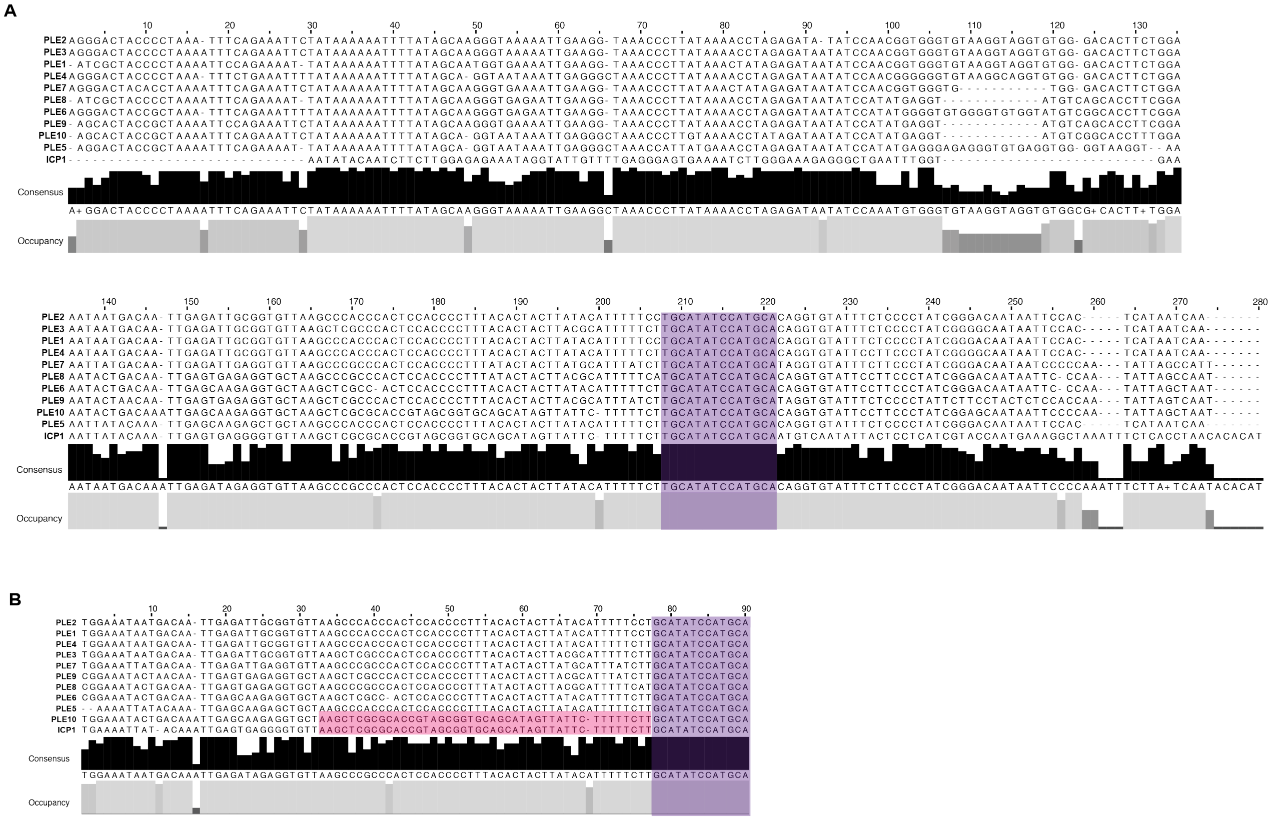

**Figure S1. The putative *pac* site is conserved in PLEs 1-10.**

- **A)** Alignment of ICP1 and PLE *pac* site regions (~250 base pairs). The 100% conserved 14-base pair region is highlighted in purple.
- **B)** Alignment of ICP1 and PLE *pac* site regions (~90 base pairs). The 100% conserved 14-base pair region is highlighted in purple, and the region of PLE10 showing more similarity to ICP1 is highlighted in pink.

**A**

| Orf3 | PLE3 | PLE4 | PLE6 | PLE10 | PLE7 | PLE8 | PLE2 | PLE1 | PLE5 | PLE9 |
|---|---|---|---|---|---|---|---|---|---|---|
| PLE3 | | 99.24 | 98.47 | 98.47 | 96.18 | 96.18 | 95.42 | 19.58 | 18.88 | 21.68 |
| PLE4 | 99.24 | | 99.24 | 99.24 | 95.42 | 95.42 | 94.66 | 20.28 | 19.58 | 22.38 |
| PLE6 | 98.47 | 99.24 | | 99.24 | 95.42 | 95.42 | 94.66 | 20.28 | 19.58 | 22.38 |
| PLE10 | 98.47 | 99.24 | 99.24 | | 95.42 | 95.42 | 94.66 | 20.28 | 19.58 | 22.38 |
| PLE7 | 96.18 | 95.42 | 95.42 | 95.42 | | 100.00 | 99.24 | 18.18 | 17.48 | 20.28 |
| PLE8 | 96.18 | 95.42 | 95.42 | 95.42 | 100.00 | | 99.24 | 18.18 | 17.48 | 20.28 |
| PLE2 | 95.42 | 94.66 | 94.66 | 94.66 | 99.24 | 99.24 | | 17.48 | 16.78 | 19.58 |
| PLE1 | 19.58 | 20.28 | 20.28 | 20.28 | 18.18 | 18.18 | 17.48 | | 97.73 | 93.94 |
| PLE5 | 18.88 | 19.58 | 19.58 | 19.58 | 17.48 | 17.48 | 16.78 | 97.73 | | 94.70 |
| PLE9 | 21.68 | 22.38 | 22.38 | 22.38 | 20.28 | 20.28 | 19.58 | 93.94 | 94.70 | |

**B**

| Orf4 | PLE7 | PLE3 | PLE4 | PLE6 | PLE2 | PLE5 | PLE9 | PLE1 | PLE10 | PLE4 |
|---|---|---|---|---|---|---|---|---|---|---|
| PLE7 | | 98.39 | 96.83 | 96.83 | 95.28 | 19.38 | 19.38 | 19.38 | 30.47 | 28.91 |
| PLE3 | 98.39 | | 98.40 | 98.40 | 96.83 | 19.53 | 19.53 | 19.53 | 30.71 | 29.13 |
| PLE4 | 96.83 | 98.40 | | 100.00 | 98.41 | 19.23 | 19.23 | 19.23 | 30.23 | 28.68 |
| PLE6 | 96.83 | 98.40 | 100.00 | | 98.41 | 19.23 | 19.23 | 19.23 | 30.23 | 28.68 |
| PLE2 | 95.28 | 96.83 | 98.41 | 98.41 | | 19.08 | 19.08 | 19.08 | 30.00 | 28.46 |
| PLE5 | 19.38 | 19.53 | 19.23 | 19.23 | 19.08 | | 100.00 | 97.62 | 86.61 | 85.83 |
| PLE9 | 19.38 | 19.53 | 19.23 | 19.23 | 19.08 | 100.00 | | 97.62 | 86.61 | 85.83 |
| PLE1 | 19.38 | 19.53 | 19.23 | 19.23 | 19.08 | 97.62 | 97.62 | | 84.25 | 83.46 |
| PLE10 | 30.47 | 30.71 | 30.23 | 30.23 | 30.00 | 86.61 | 86.61 | 84.25 | | 98.41 |
| PLE8 | 28.91 | 29.13 | 28.68 | 28.68 | 28.46 | 85.83 | 85.83 | 83.46 | 98.41 | |

**C**

| Orf5 | PLE7 | PLE3 | PLE4 | PLE2 | PLE6 | PLE1 | PLE9 | PLE8 | PLE5 | PLE10 |
|---|---|---|---|---|---|---|---|---|---|---|
| PLE7 | | 98.96 | 97.92 | 97.92 | 97.92 | 34.38 | 34.38 | 36.46 | 36.46 | 36.46 |
| PLE3 | 98.96 | | 98.96 | 98.96 | 98.96 | 34.38 | 34.38 | 36.46 | 36.46 | 36.46 |
| PLE4 | 97.92 | 98.96 | | 100.00 | 98.96 | 34.38 | 34.38 | 36.46 | 36.46 | 36.46 |
| PLE2 | 97.92 | 98.96 | 100.00 | | 98.96 | 34.38 | 34.38 | 36.46 | 36.46 | 36.46 |
| PLE6 | 97.92 | 98.96 | 98.96 | 98.96 | | 34.38 | 34.38 | 36.46 | 36.46 | 36.46 |
| PLE1 | 34.38 | 34.38 | 34.38 | 34.38 | 34.38 | | 97.83 | 95.65 | 92.39 | 92.39 |
| PLE9 | 34.38 | 34.38 | 34.38 | 34.38 | 34.38 | 97.83 | | 95.65 | 92.39 | 92.39 |
| PLE8 | 36.46 | 36.46 | 36.46 | 36.46 | 36.46 | 95.65 | 95.65 | | 90.22 | 90.22 |
| PLE5 | 36.46 | 36.46 | 36.46 | 36.46 | 36.46 | 92.39 | 92.39 | 90.22 | | 100.00 |
| PLE10 | 36.46 | 36.46 | 36.46 | 36.46 | 36.46 | 92.39 | 92.39 | 90.22 | 100.00 | |

**Figure S2. Comparisons of Orf3, Orf4, and Orf5 protein variants.**

A) Pairwise comparison of percent identity of each Orf3 protein from 10 PLEs.
B) Pairwise comparison of percent identity of each Orf4 protein from 10 PLEs.
C) Pairwise comparison of percent identity of each Orf5 protein from 10 PLEs.

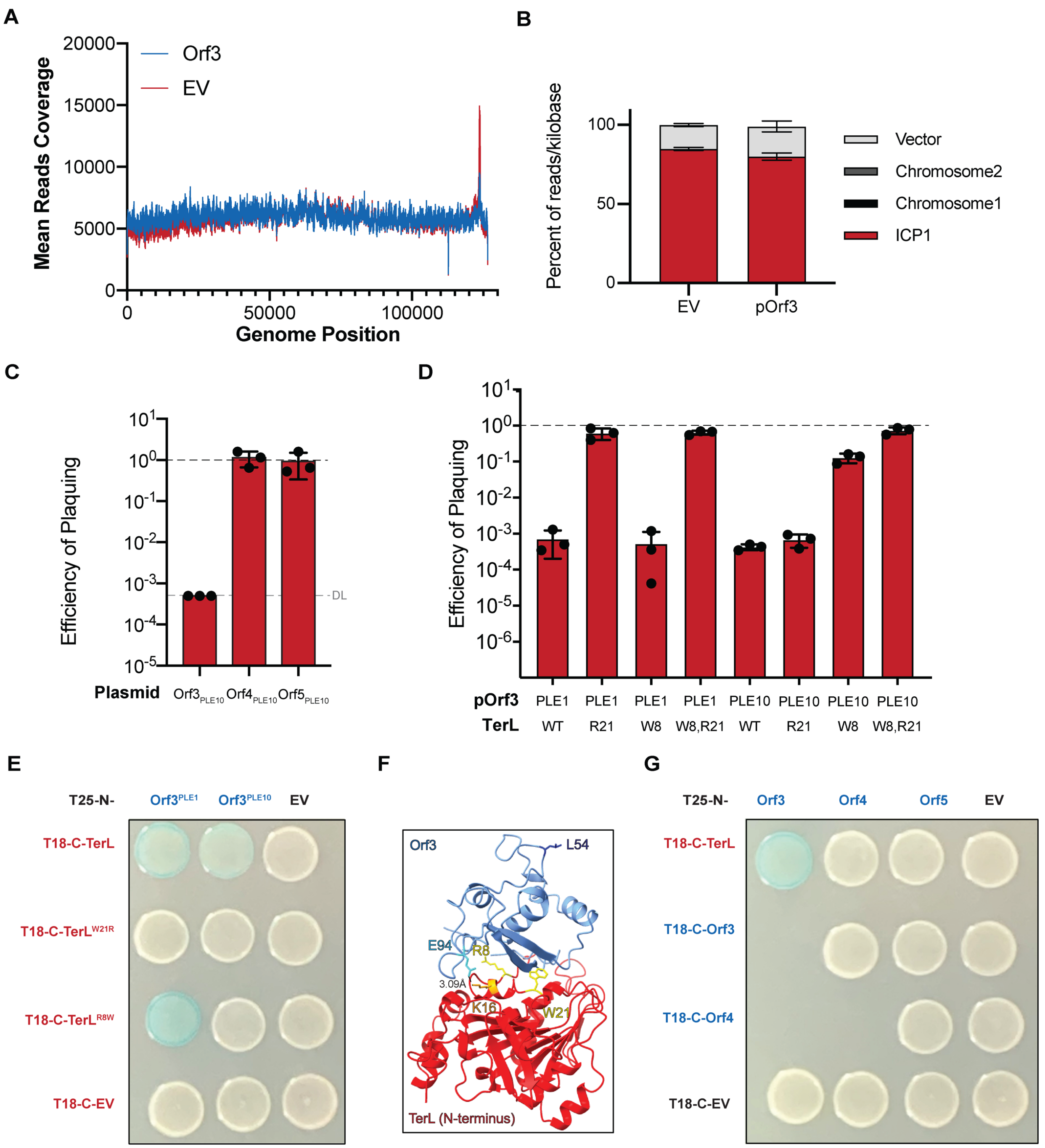


**Figure S3. The two variants of Orf3 function similarly to bind and inhibit ICP1's TerL.**

A) Mean reads coverage of ICP1's genome late during infection of *V. cholerae* carrying an empty vector (EV) or a plasmid expressing $Orf3_{PLE1}$. The peak corresponds to redundantly packaged sequence around the predicted *pac* site where reads are markedly lower for the Orf3-expressing strain compared to the strain with the empty vector (see Figure 2D for a zoomed-in of this region).

B) Average percent distribution of mapped reads/kilobase for the four genetic elements in the cell late during infection. Error bars show standard deviation (n=3).

C) ICP1 efficiency of plaquing on strains expressing alleles from PLE10 relative to an empty vector. The gray dashed line represents the detection limit.

D) Efficiency of plaquing of ICP1 harboring TerL substitutions on strains expressing $Orf3_{PLE1}$ and $Orf3_{PLE10}$.
E) Representative BACTH results for $Orf3_{PLE1}$ and $Orf3_{PLE10}$ with TerL, $TerL^{W21R}$, and $TerL^{R8W}$ (n=3).
F) Alpha-Fold2 predicted structures and interactions for $Orf3_{PLE10}$ (light blue) and the N-terminus of TerL (red). Residues that are substituted to escape $Orf3_{PLE1}$ (W21) and $Orf3_{PLE10}$ (R8) are shown in yellow, residue for the $Orf3_{PLE1}$ variant that is predicted to interact with TerL (L54) is in dark blue, and the predicted interaction between $Orf3_{PLE10}$ E94 (cyan) and TerL K16 (orange) is within 4Å.
G) Representative BACTH results for $Orf3_{PLE1}$, $Orf4_{PLE1}$, and $Orf5_{PLE1}$ and TerL (n=3).

For C and D, the black dashed line indicates an efficiency of 1 equal to the empty vector control. Each dot represents a biological replicate, and the error bars indicate the mean between the replicates (n=3).
